# Hydrogen and dark oxygen drive microbial productivity in diverse groundwater ecosystems

**DOI:** 10.1101/2022.08.09.503387

**Authors:** S. Emil Ruff, Pauline Humez, Isabella Hrabe de Angelis, Michael Nightingale, Muhe Diao, Sara Cho, Liam Connors, Olukayode O. Kuloyo, Alan Seltzer, Samuel Bowman, Scott D. Wankel, Cynthia N. McClain, Bernhard Mayer, Marc Strous

## Abstract

Groundwater ecosystems are globally wide-spread yet still poorly understood. We investigated the age, aqueous geochemistry, and microbiology of 138 groundwater samples from 87 monitoring wells (<250m depth) located in 14 aquifers in the Canadian Prairie. Geochemistry and microbial ecology were tightly linked revealing large-scale aerobic and anaerobic hydrogen, methane, nitrogen, and sulfur cycling carried out by diverse microbial communities. Older groundwaters contained on average more cells (up to 1.4×10^7^/mL) than younger groundwaters. Organic carbon-rich strata featured some of the highest abundances, challenging current estimates of global groundwater population sizes. Substantial concentrations of dissolved oxygen (n=57; 0.52±0.12 mg/L [mean±SE]; 0.39 mg/L [median]) in older groundwaters could support aerobic lifestyles in subsurface ecosystems at an unprecedented scale. Metagenomics, oxygen isotope analyses and mixing models indicated that microbial “dark oxygen” contributed to the dissolved oxygen pool in subsurface ecosystems commonly assumed to be anoxic.

---

About 2 % of Earth’s water resources occur as groundwater, of which half is saline and the other half fresh (*1*). This subsurface freshwater represents ~30 % of global freshwater resources, around sixty times more than in all lakes, rivers and the atmosphere combined, and is exceeded only by the inaccessible and currently still frozen polar ice caps (*1*). Aquifers and rock fractures may also hold up to 30 % of the total microbial biomass on Earth (*2*, *3*), contribute substantial carbon fixation (*4*), and contain high proportions of uncultured archaea, bacteria, and viruses (*2*, *5*) with a broad spectrum of lifestyles (*6*). Despite the global occurrence of groundwater and the magnitude and diversity of its resident biomass, our understanding of the composition and activity of microbial communities inhabiting these hidden aquatic ecosystems is still patchy, often developed from samples of a few select wells or a single aquifer. In particular, the geochemical and ecological processes that shape groundwater microbial communities over space and time are not well constrained (*7*).

Establishing robust links between microbial communities and geochemical characteristics of groundwaters requires large datasets having a comprehensive environmental inventtory associated with each microbial community sample. To this end, the Groundwater Observation Well Network (GOWN) maintained by Alberta Environment and Parks (AEP) in Canada, has compiled geochemical data for over 250 groundwater samples obtained from monitoring wells in different aquifers and geographic regions, representing a variety of geochemical regimes and groundwater ages. Each GOWN well has been sampled repeatedly for many years, including some for several decades (*8*). Since 2006, this comprehensive monitoring program has systematically collected regular water level and chemical and isotopic water quality information for aqueous and gaseous samples (*9*).

The province of Alberta is situated in the Western Canadian Sedimentary Basin, which hosts major oil, gas, coal, as well as sulfur, salt, limestone, and dolomite deposits (*10*). The shallow and deep subsurface has been extensively studied in the context of petroleum and coal exploration and development (*11*)(Fig 1). Here, we analyzed 138 groundwater samples from 87 GOWN wells completed in bedrock and surficial aquifers across Alberta (Fig 1, 1) with the aim of investigating the processes that govern microbial community ecology in freshwater aquifers. To achieve this aim, we employed a multidisciplinary characterization of the groundwater geochemical composition including dissolved gases, determination of groundwater ages and the composition of its resident microbial communities. The guiding objectives were to determine (i) whether geochemistry is linked to microbial community structure, (ii) which energy sources support these communities and (iii) which microbial lineages represent key populations.

**Fig. 1.**
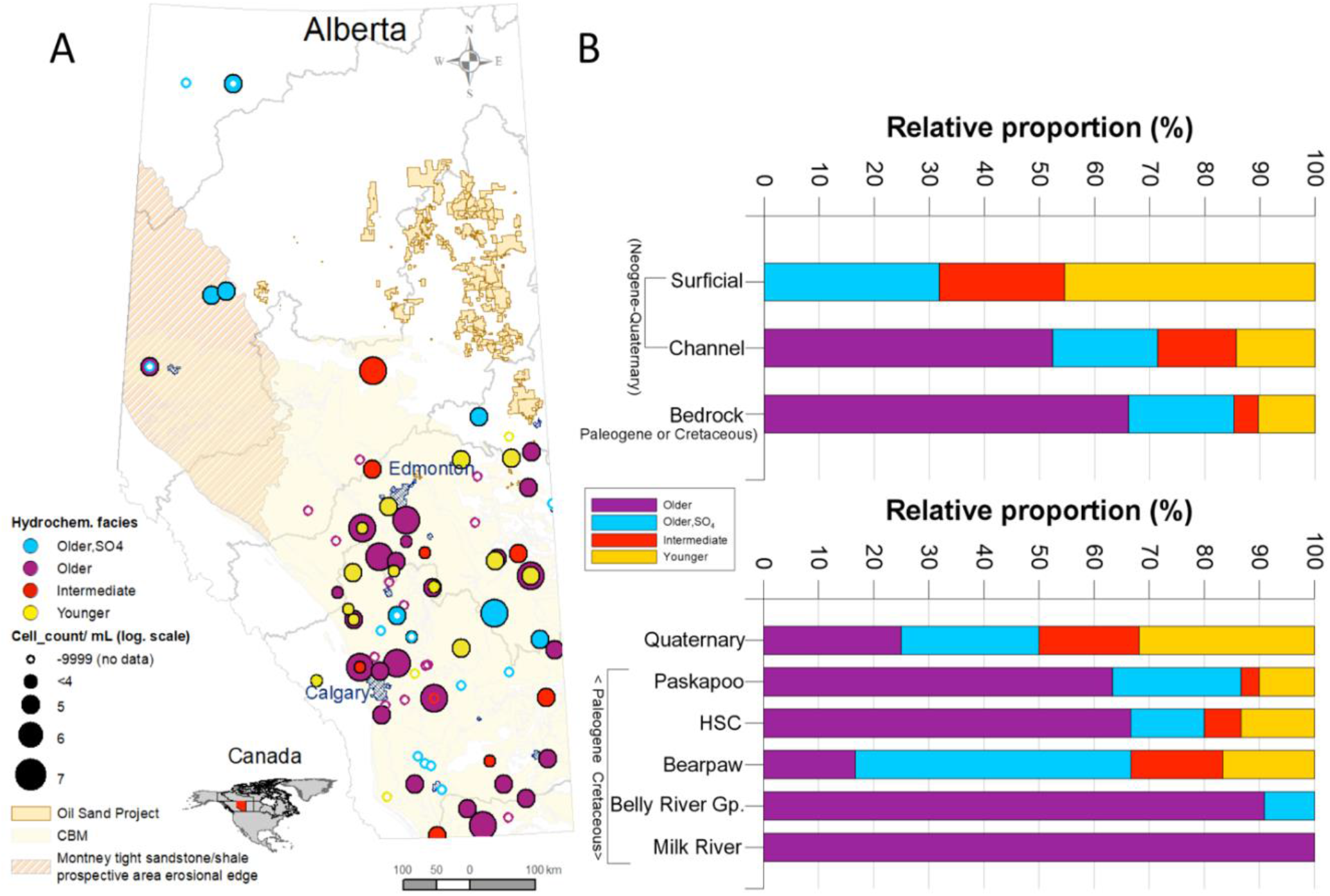
Sampling locations, geological formations, and groundwater ages. **(A**) Location of studied groundwater wells within the energy resources context of the province of Alberta. Colors indicate the groundwater age at each well (yellow: younger waters; red: intermediate age; purple: older waters; blue: older waters sulfate-rich). Circle size represents average microbial cell numbers in the groundwater samples, ranging from 10^4^ (smallest full circle) to 10^7^ cells per mL (largest circle). CBM: Coal-bed methane. B) Relative proportion of water types in the surficial, channel and bedrock sediments, as well as in major geological formations of Alberta, showing that groundwater geochemistry evolved with increasing age of the formations. HSC: Horseshoe canyon.

## Geochemical evolution of groundwater in the Canadian Prairies

The mean residence time of water in an aquifer is a very important factor influencing its geochemical composition (or facies) (Fig 2A). Mean residence time, or groundwater age, is determined using radioisotopes such as tritium and ^14^C and refers to the travel time between the point of infiltration and the point of sampling (*12*). In this study, groundwater age inversely correlated with the ratio between calcium and sodium concentrations in groundwater (Fig 2B). The youngest, tritium-containing groundwaters (indicating recharge after 1960) were characterized by low total dissolved solids (< 400 mg/L) and high Ca/Na ratios (median: 3.5).

**Fig. 2.**
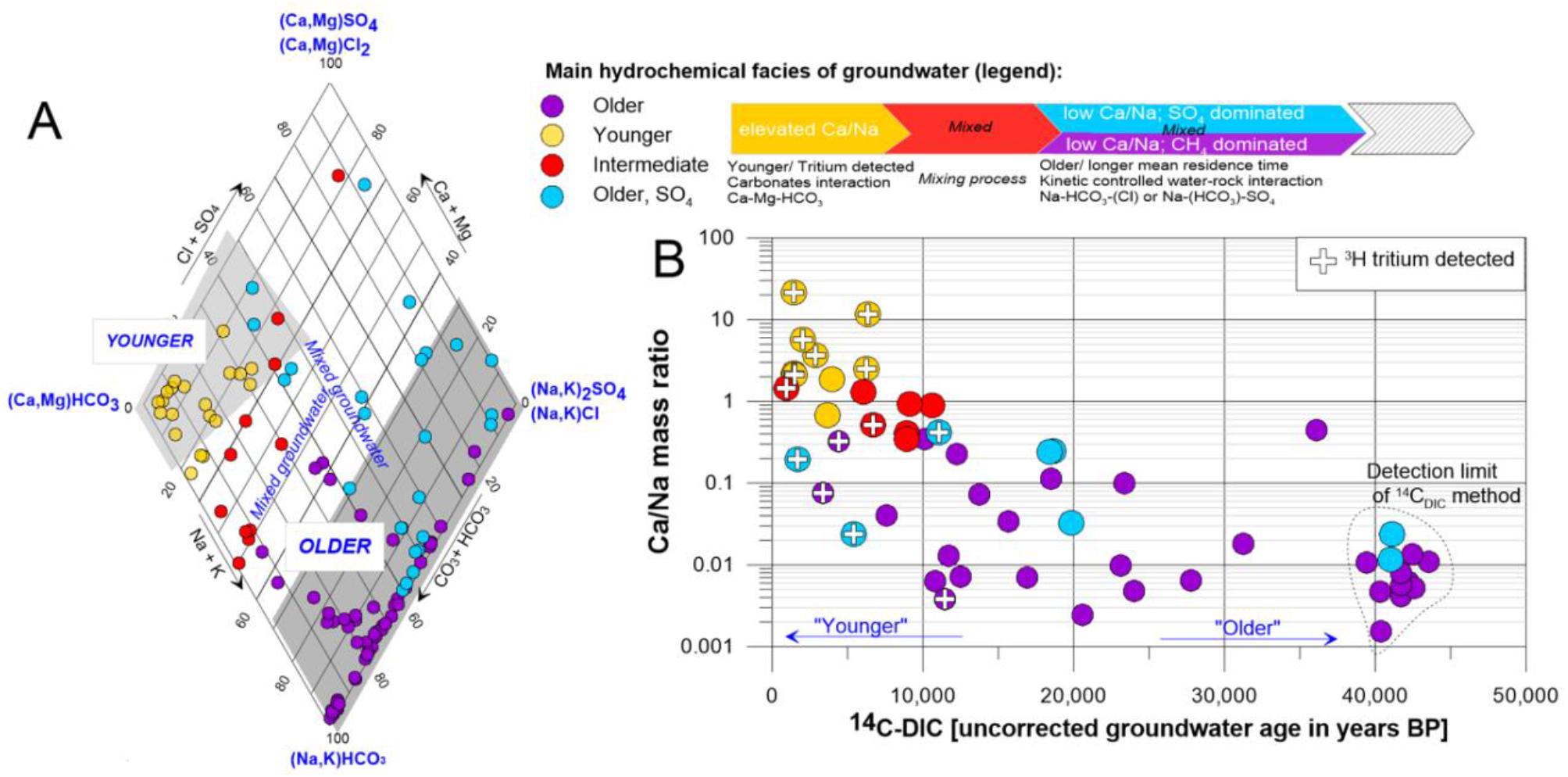
Groundwater geochemistry and age dating. (**A**) Piper diagram of hydrochemical facies of each groundwater sample (circles) visualizing calcium/sodium mass ratio used as a proxy for geochemical evolution, i.e., residence time/age. Schematic timeline of water aging. (**B**) Ca/Na ratio decreases with increasing residence time (^14^C_DIC_ uncorrected age). ^3^H-positive samples (circles with crosses) corroborate the age trend.

Dissolution of carbonates during infiltration led to high calcium, magnesium, and bicarbonate concen-trations in these waters. These young ground-water samples (yellow symbols, Fig 2) generally had low methane and sulfate concentrations (Fig 3A, B) and were pre-dominantly collected from wells completed in surficial Neogene-Quaternary deposits (Fig 1B).

**Fig. 3.**
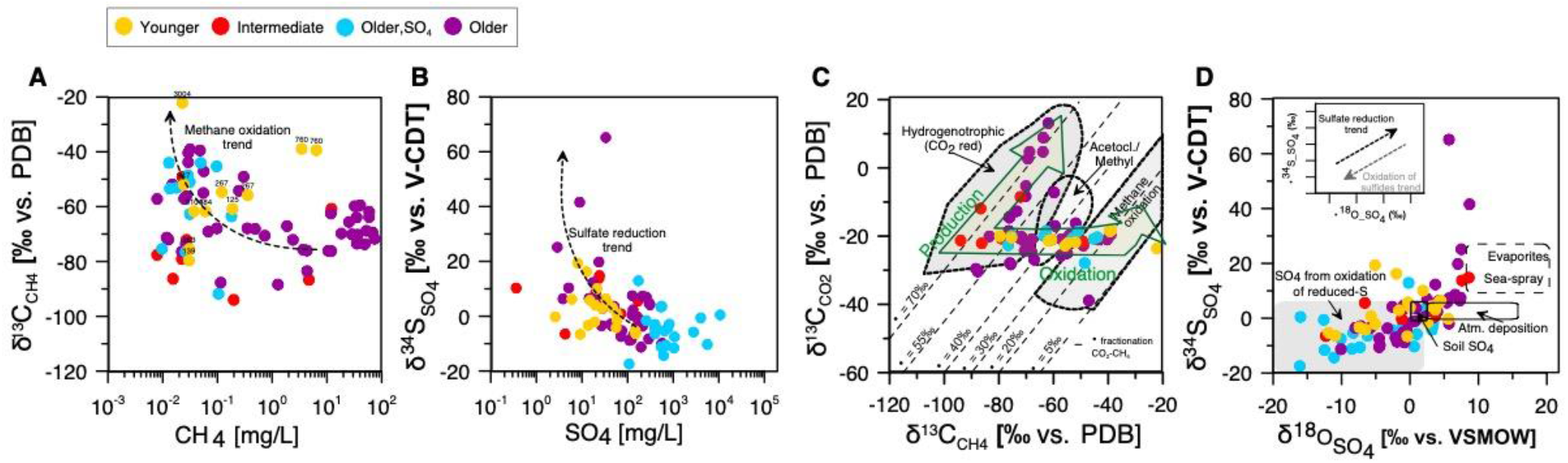
Groundwater isotope geochemistry. Microbes mediate methane (CH_4_) and sulfate (SO_4_^2-^) cycling impacting carbon and sulfur pools in the groundwater systems. (**A**) Carbon isotopic signature of methane versus methane concentration. (**B**) Sulfur isotopic signature of sulfate versus sulfate concentration. (**C**) Carbon isotopic signature of carbon dioxide versus carbon isotopic signature of methane. (**D**) Sulfur isotopic signature of sulfate versus oxygen isotopic signature of sulfate. Each circle is a sample. Circle color depicts water age. Trends concerning reduction and oxidation of the compounds are indicated by arrows or areas.

In contrast, the oldest groundwaters (purple symbols, Fig 2) contained no tritium and little ^14^C, indicative of groundwater more than several hundreds or thousands of years old. They had elevated total dissolved solids (> 900 mg/L) and a low Ca/Na ratio (median: 0.01). Old groundwaters were characterized by reducing conditions and contained high dissolved methane concentrations (12.8±2.4 mg/L [mean±SE]; median: 0.72 mg/L, range: 0.001-74.2 mg/L; Fig 3A, Fig S1). They had elevated sodium, bicarbonate, and chloride concentrations resulting from water-rock interactions, including ion exchange, and weathering of minerals. The older groundwater samples were obtained from wells completed in buried river valleys (channels) and Paleogene and Cretaceous sedimentary bedrock formations that are often characterized by the presence of coal and/or shale (*13*).

Sulfate, predominantly derived from pyrite oxidation (Fig 3D) and to a lesser extent from anhydrite or gypsum dissolution (*14*), was ubiquitous in a third group of groundwater samples (blue symbols, Fig 2). These waters were characterized by long residence times, contained even higher dissolved solids (> 1700 mg/L) and had intermediate Ca/Na (median of 0.12). Sulfate was often the most abundant anion and electron acceptor in this group of groundwaters, resulting in a sulfate-rich hydrochemical facies with low methane concentrations. These groundwater samples were collected from wells completed in surficial deposits, but also from bedrock aquifers completed in clastic, often marine sedimentary rocks of the Bearpaw formation (Fig 1B).

The mixing of groundwater of different ages and geochemical water types (Fig 2A) often resulted in characteristic intermediate hydro-chemical facies (red symbols, Fig 2). These waters were mostly obtained from aquifers in surficial deposits and occasionally from bedrock aquifers (Fig 1B). A detailed characterization of the geologic formations, groundwater geochemistry and ages is provided in the Supplementary Materials and Data S1.

## Geochemically evolved groundwater contains biomass-rich microbial communities

Microbial cell density commonly decreases with increasing depth in marine (*15*) and terrestrial (*3*) subsurface ecosystems. This decrease in cell numbers and biomass is often attributed to energy limitation in subsurface ecosystems (*16*). Within aquifers cell numbers may not necessarily decrease with increasing depth (*17*), but due to other factors including increasing water age (*18*, *19*). To our surprise, we found significantly more cells in aquifers with old groundwater than in those with younger water (Fig 4). This indicated that the sampled aquifers are in fact productive ecosystems that provide energy for growth of microorganisms not only in shallow and young groundwater but also in aquifers with old and geochemically evolved groundwater.

**Fig. 4.**
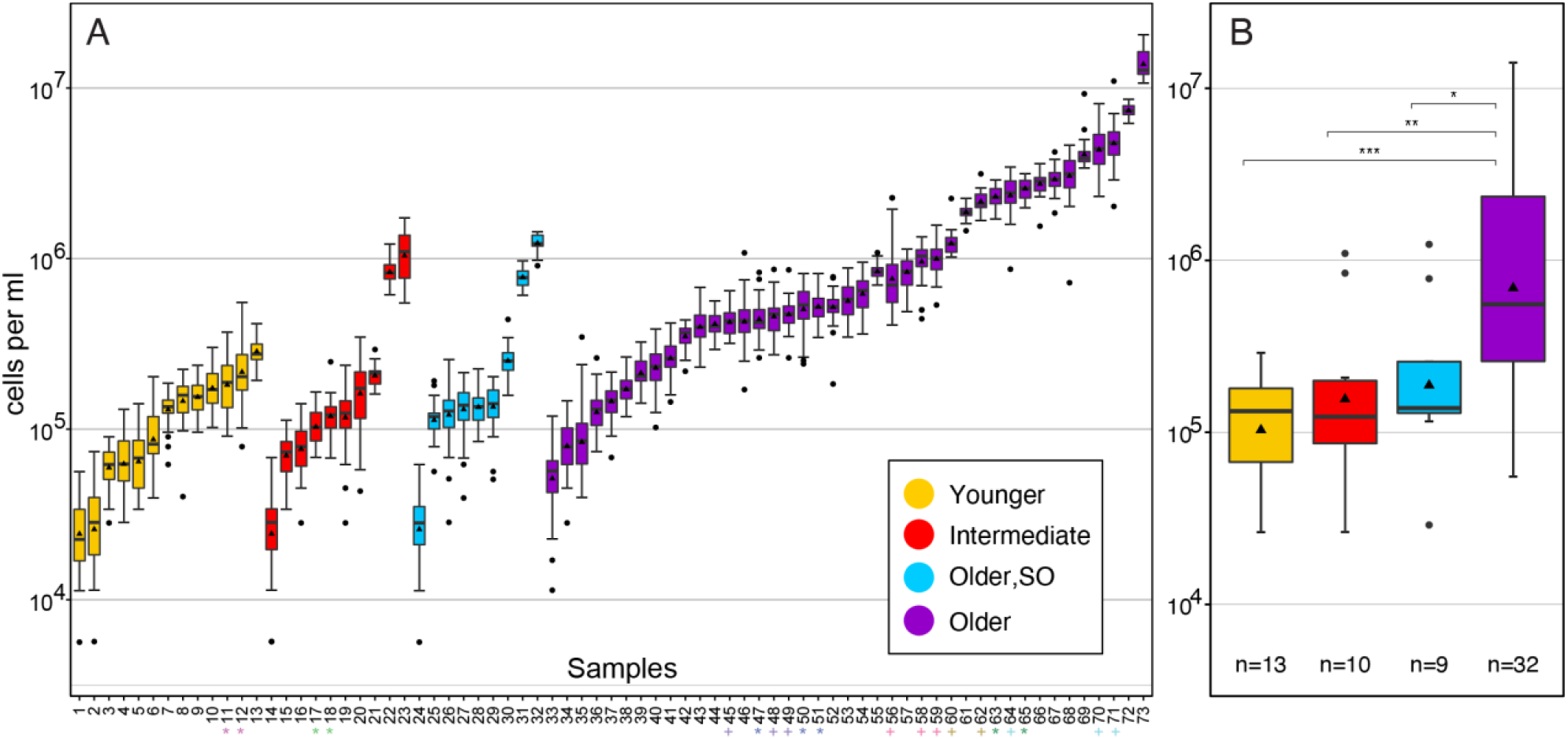
Cell abundances in groundwaters. Cell abundance was determined using fluorescence microscopy and is given on a logarithmic scale. Boxplots show 1^st^ and 3^rd^ quartile, maximum and minimum values (whiskers), median (line), mean (triangle), and outlier (dots). (**A**) Boxplots represent groundwater samples from individual wells and summarize cell counts from 20-40 independent fields of view (details see Data S1). Wells which were sampled at two different time points are marked with a star (*), technical replicates are marked with a plus (+). (**B**) Boxplots represent cell numbers averaged across groundwater age summarizing the average cell numbers in panel A (without technical replicates). n: Number of samples. Significance was tested using a Wilcoxon rank sum test. Significance levels are: *: p<0.05; **: p<0.01; ***: p<0.001.

Average cell numbers in the geochemically evolved groundwater samples reached 10^7^ cells per ml (Fig 4). Overall, microbial cell numbers across the entire studied region, did not decrease with depth (Fig S2A, B). Cell counts were on average slightly higher in aquifers in geologic strata that contained shales or coal (Fig S2C), suggesting that occurrence of elevated contents of organic carbon may provide additional energy sources to microbes. It should be noted that the cell numbers presented here are conservative estimates capturing only the free-living cells from water samples which cannot account for the substantial number of cells living in biofilms (*20*).

The morphology and size of cells varied greatly, and included cocci-, rod-, vibrio-, and spiral-shaped cells, as well as filaments and small aggregates (Fig S3, Data S2). Cell morphologies, sharp cell boundaries and the bright signal from nucleic acid staining suggested a largely active community (*21*). Average cell size was similar across groundwater samples and hence high cell numbers are a proxy for high biomass and productivity (Fig S3, Data S2). To the best of our knowledge this is the first time that very high levels of biomass were documented in aquifers across a large geographic area (>210,000 km^2^).

The increase in cell numbers with groundwater age was accompanied by a substantial decrease in archaeal and bacterial diversity (Fig 5A, B) and a shift in microbial community structure from young to old groundwater (Fig S4, S5). The environmental parameters most strongly associated with variance in microbial community structure were methane concentrations and geochemical proxies for groundwater age, including the concentrations of sodium, calcium, and magnesium and well depth (Fig 5C, D). Decreases in diversity and shifts in community structure are common features of microbial bloom situations and highly productive ecosystems, which often feature elevated abundance of a few select species (*22*).

**Fig. 5.**
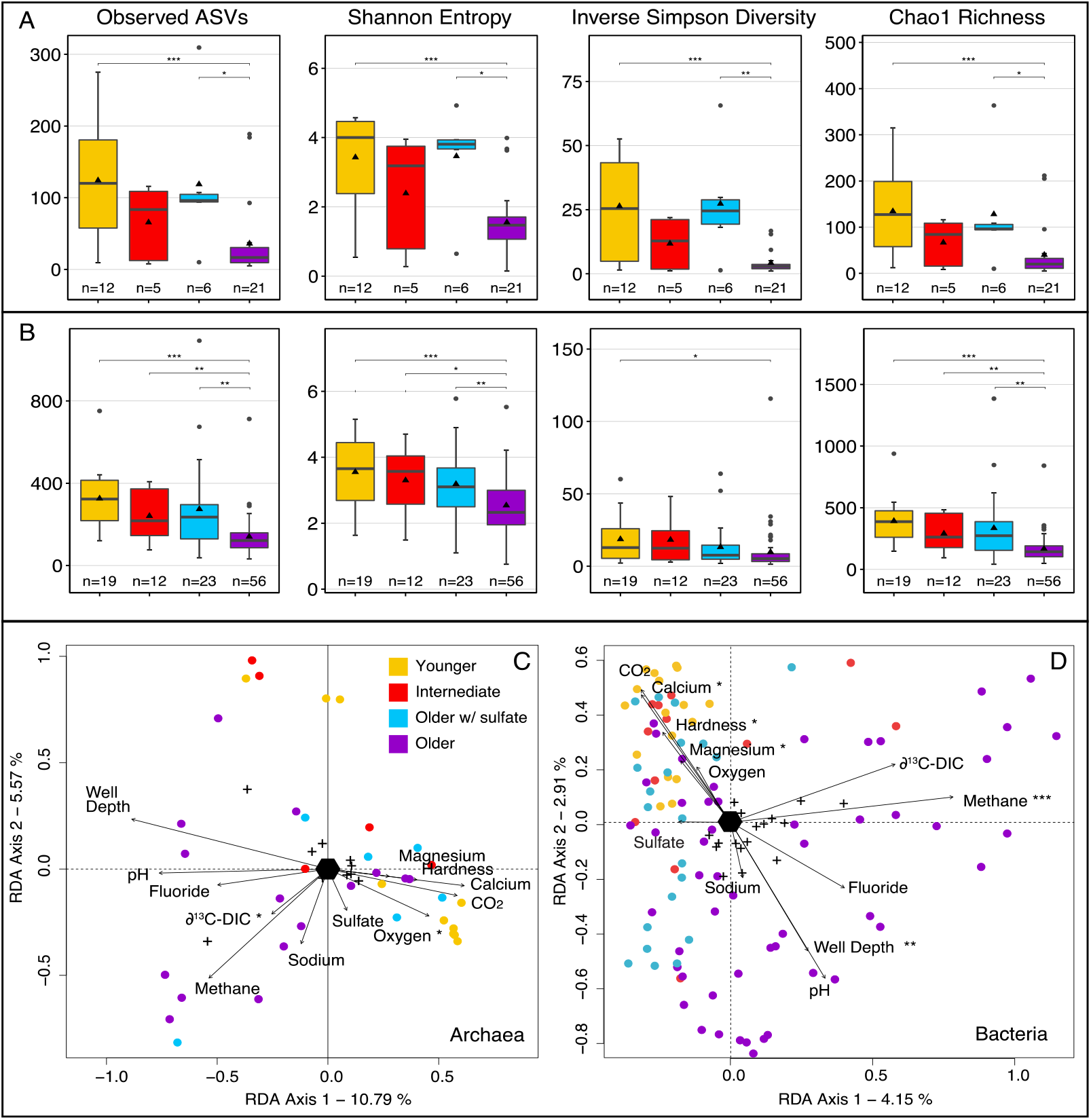
Microbial community diversity. Archaeal (**A**) and bacterial (**B**) alpha diversity indices of the investigated groundwaters based on 16S rRNA gene amplicon sequence variants (ASVs). Significance was tested using a Wilcoxon rank test. Significance levels are: *: p<0.05; **: p<0.01; ***: p<0.001. Redundancy analysis of the archaeal (**C**) and bacterial (**D**) community structure in groundwater samples (circles) using the most important parameters (arrows) as determined by EnvFit. Significance levels: ***: p<0.001, **: p<0.01, *: p<0.05. The full model was highly significant for both domains, and together the eleven parameters explained 24 % of archaeal and 18 % of bacterial variation.

## Hydrogen as basal energy source in the high productivity aquifers

Our results indicated that a major energy source driving high productivity in mature groundwater is hydrogen. The detection of considerable amounts of hydrogen was shown only at selected sites (0.002, 0.007 and 0.1 % in samples GW121, GW223 and GW951), yet in many aquifers obligate hydrogenotrophic methanogens including *Methanobacterium, Methanoregula* and *Methanospirillum* showed high relative abundances of 16S rRNA gene sequences (Fig 6A, S7, Data S3). Hydrogen serves as sole electron donor for these lineages and widespread hydrogeno-trophic methanogenesis is also sup-ported by the isotope composition of CH_4_ and CO_2_ (Fig 3C). Methane with concentrations ranging from 0.006 to more than 74 mg/L was present in all analyzed groundwater samples (Fig 3A). Most samples, however, had concentrations <1 mg/L (73 %; n=106, Fig S6). Based on combined carbon and hydrogen isotope composition, these low methane concentrations likely reflect high relative consumption by microbial methane oxidation (Fig 3A).

**Fig. 6.**
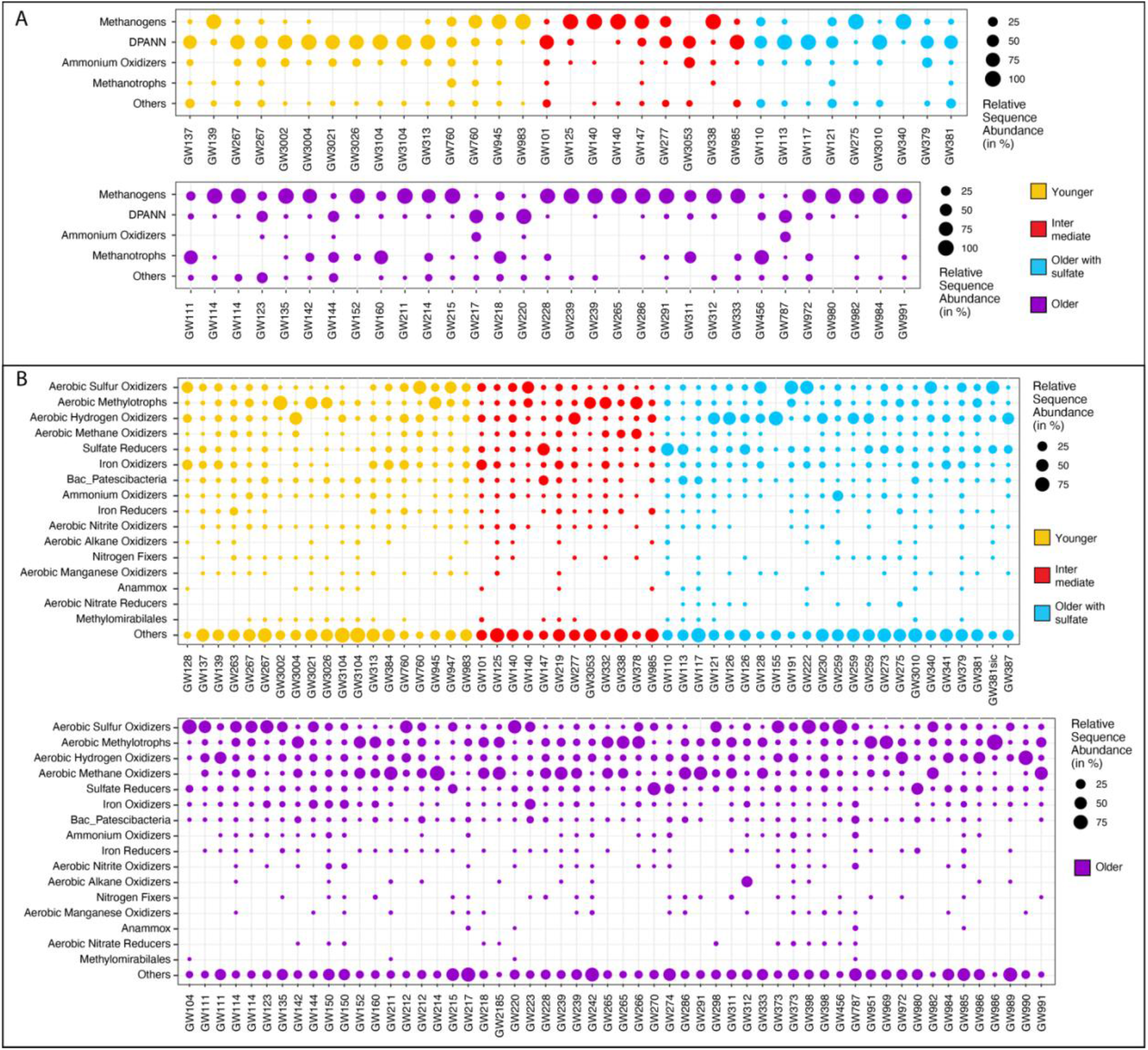
Microbial community composition. Relative sequence abundance of important archaeal (A) and bacterial (B) guilds. Together these clades are likely responsible for the majority of element cycling that was detected. The members of each guild were selected based on taxonomic information affiliated with each sequence, e.g., sequences affiliating with the genus *Methanobacterium* were assumed to represent obligate, hydrogenotrophic methanogens. We avoided to include known mixotrophs, hence the guilds are rather conservative. We are aware that inferences on metabolism and activity of an organism based on its presence in a 16S rRNA gene amplicon dataset has limitations, but in conjunction with the comprehensive geochemical and isotopic data as well as cell numbers we believe that these data represent a good approximation of the actual community functions.

Among the methane oxidizers were obligate anaerobic methane oxidizing archaea (ANME) of the *Methanoperedenaceae* (ANME-2d) often found in aquifers that contained either dissolved iron or manganese or both. ANME-2d were found in more than half of the archaeal community datasets (34 of 64), predominantly in methane-rich and sulfate-poor older groundwaters, in line with their occurrence in deep granitic water environments (*23*). Their ability to oxidize methane using iron and manganese oxides (*24*) could explain the high relative abundance of over 40 % in the sulfate depleted mature groundwaters of at least four samples (GW111, GW218, GW311 and GW456). The genus *Methanoperedens* was the fifth most abundant archaeal genus in the studied aquifers and has the metabolic potential to connect the carbon, nitrogen, and metal cycles via the anaerobic oxidation of methane (*25*,*26*). Anaerobic oxidation of methane coupled to sulfate reduction seemed to occasionally occur as well, as ANME-2ab made up almost half of the archaeal community at GW160 and ANME-2c were detected at GW220. Bacterial *Methanomirabiliales*, capable of oxidizing methane in anoxic environments using nitrite (*27*), were abundant in many aquifers, especially in young and intermediate waters (Fig S8).

Hydrogen likely also fueled growth of organisms affiliating with the genera *Desulforudis* and *Desulfomicrobium* and many other sulfate and sulfur reducers, and sulfur disproportionators (Fig S9), which were diverse and widespread in the studied groundwater eco-systems (Fig 6B, S6). *Candidatus* Desulforudis audaxviator were the most sequence abundant sulfate reducers across all samples, yet predominantly occurred in old groundwaters. They often made up more than 50 % of sulfur cycling microbes. *Ca*. Desulforudis are hydrogen-oxidizing, sulfate-reducing *Clostridia* reported to thrive in deep terrestrial aquifer eco-systems (*28*). In samples GW3026 and GW217 we found high sequence abundances of obligately syntrophic *Smithella sp*. and *Syntrophus sp*. that thrive together with organisms scavenging hydrogen (*29*, *30*). The origin of the hydrogen is unclear, yet the gas could be derived from microbial fermentation, and from abiotic sources including shales, coal beds and other organic rich strata (*31*), water-rock reactions (*32*), pumping equipment or steel well casings (*33*), or even radiolysis (*34*, *35*).

## Aquifers contained widespread and diverse aerobic microbial lineages

We found high relative sequence abundances of obligate and facultatively aerobic hydrogen-, methane-, and sulfide-oxidizing lineages in most of the old groundwaters obtained from confined aquifers (Fig 6, 7, 9, S7–10, Data S4). The presence of aerobic organisms was not the result of a contamination, or of microbial growth after sampling, because alpha and beta diversity, composition and cell counts did not change with storage time (Fig S11) and thus we are confident that these aerobes were present in the aquifers at the time of sampling. We detected high relative sequence abundances of aerobic hydrogen oxidizers affiliating with the genus *Hydrogenophaga* (Fig 6A, 7A). Obligately aerobic methano- and methylotrophic bacteria (*36*) were found in all aquifers, especially in older groundwater. These bacteria were often the most abundant bacteria in a sample (Fig 6B, S8). Aerobic methylotrophic *Methylotenera sp*. were the most abundant genus-level clade in the dataset overall, followed by aerobic methane-oxidizing members of the genus *Methylobacter*.

**Fig. 7.**
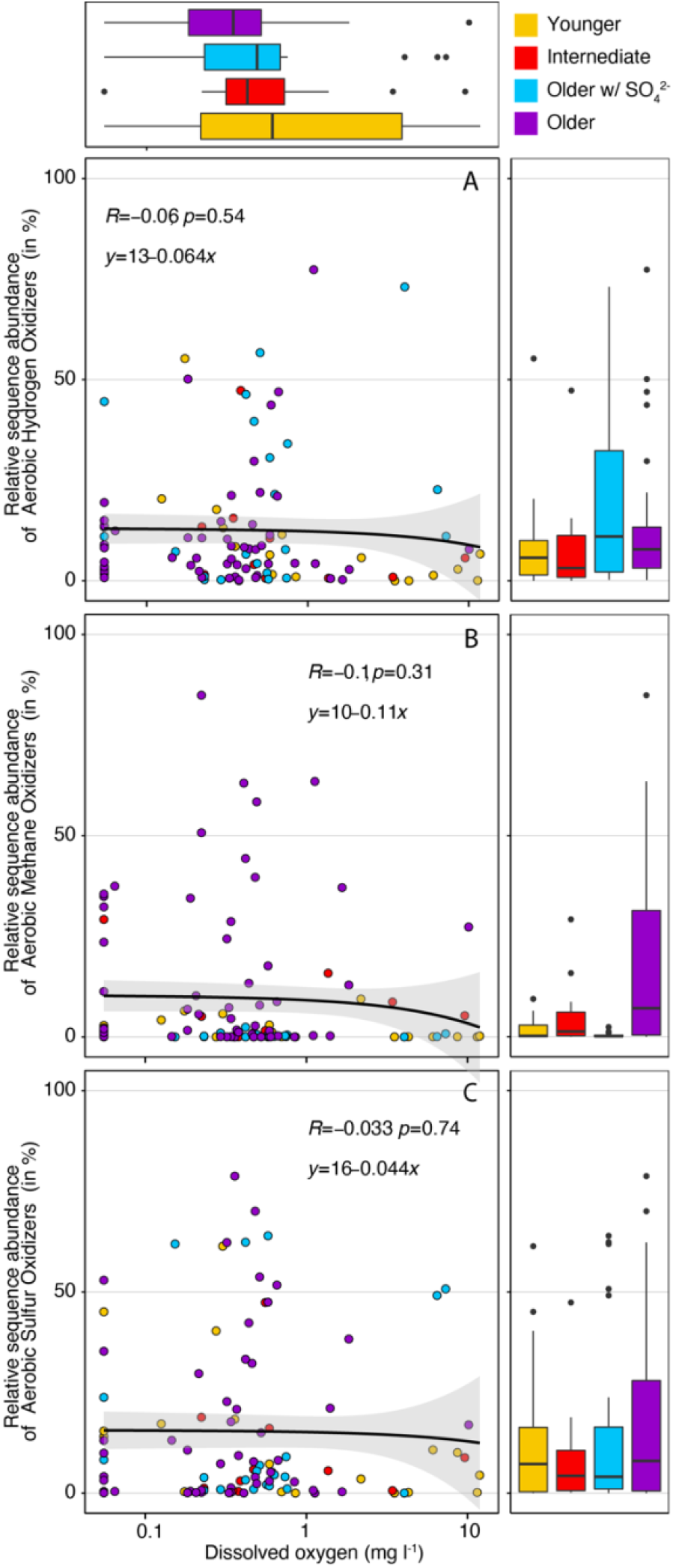
Aerobic microbial clades: Relative abundance of aerobic hydrogenotrophs (**A**), methanotrophs (**B**) and thiotrophs (**C**) versus dissolved oxygen concentration. Relative sequence abundance of the aerobic clades is always highest at intermediate oxygen concentrations indicating that these waters contain both sufficient electron donors (hydrogen, methane, sulfur) and electron acceptor (oxygen). Oxygen concentration is shown on pseudo-log10 scale, to include the samples in which oxygen was not detected (zero). Boxplots summarize relative abundances (right panels) and oxygen concentrations (top panel) using water age.

The coexistence of these guilds was previously observed in a methane-rich shallow aquifer (*37*) as well as in methylotrophic microbiomes of mesocosms (*38*). Aerobic methane oxidizers of the genus *Crenothrix* (*39*) were the fourth most abundant and occurred in almost half of the samples (45 of 109, Fig S8). *Crenothrix sp*. are filamentous microbes that can occur in groundwater (*40*) and may represent the filamentous cells observed with the microscope in many samples (Fig S3).

Of the twenty most abundant bacterial genera known to oxidize one-carbon compounds that we found in the groundwaters (Fig S8), twelve were associated with oxidation of methanol, methylamines, and other methyl compounds (*41*), while eight were associated with methane oxidation (*42*). The observed widespread signs of aerobic methanotrophy corroborates previous findings of aerobic methanotrophic activity in coalbed formations (*43*, *44*), with our study expanding this finding to more diverse geological settings and a much larger geographical scale. Facultatively anaerobic sulfide and sulfur-oxidizing bacteria including *Sulfuricurvum sp., Thiobacillus sp., Thiomicrorhabdu*s and *Sulfurimonas* were wide-spread and abundant in the studied aquifers (Fig 6C, S10). Their versatile sulfur metabolisms could explain previous observations of pyrite oxidation (*14*), as well as the presence of sulfur reducers like *Desulfuromonas* which can reduce elemental sulfur (*45*) and replenish the sulfide pool.

The biomass produced by the diverse, coexisting autotrophs in turn supports multitudes of archaeal and bacterial heterotrophs, including DPANN archaea and *Patescibacteria* (also known as Candidate Phyla Radiation). In the archaeal community aerobic *Woesearchaeales* were particularly diverse and abundant in all aquifers except those within old groundwaters, and in the bacterial community potentially heterotrophic *Comamonadaceae* and *Pseudomonadaceae* were also very abundant (Fig S12). Heterotrophic clades remineralize organic carbon compounds to carbon dioxide, which can then be recycled by hydrogenotrophic methanogens and other autotrophs to produce biomass.

## Dark oxygen is an electron acceptor in aquifers

As water infiltrates through recharge zones and flows through the aquifers, the dissolved oxygen (DO; assumed to be at saturation equilibrium initially) is consumed by microbial respiration, decomposition of organic matter, or by reacting with reduced minerals (*46*). These oxygen-consuming reactions often reduce the DO content of groundwater to below the detection limits, particularly in groundwater with long residence times that has been out of contact with the atmosphere for many years, centuries, or even millennia (*46*, *47*). To our surprise we detected low concentrations of dissolved oxygen (DO) in most groundwater samples including those of deeper aquifers containing geochemically mature groundwater (Fig 8A). Based on known stoichiometry of relevant microbial metabolisms, consumption of “missing” oxygen (DO saturation at infiltration – DO measured in the obtained samples) in 85 % of aquifers (59 of 70) could theoretically sustain more microbial cells than we observed (Data S1). In the remaining 11 aquifers, the DO content was below the detection limit. The occurrence of more than 0.3 mg/L DO in many old groundwaters of deeper and confined aquifers was remarkable and suggests that oxygen is present in ecosystems that are often assumed to be anoxic (*46*, *48*). To investigate whether the oxygen may have been introduced during sampling we carried out oxygen isotope analyses of oxygen (O_2_). Indeed, some oxygen isotope data for DO were consistent with groundwater that is in equilibri-um with atmospheric oxygen (δ^18^O_O2_ = +24 ‰ ± 0.1‰, Fig. 8B), indicating air contamination during sampling, or the presence of atmospheric-derived oxygen in the aquifers. However, at certain sites (GW218, GW265) we observed markedly lower δ^18^O_O2_ values (as low as +21 ‰), while also finding elevated O_2_:Ar ratios (Fig. 8B). The lower δ^18^O_O2_ and higher O_2_:Ar ratios interestingly fell along a trend, consistent with the simulated addition of DO with a δ^18^O value much lower than that of air-equilibrated water (dashed trend line in Fig 8B). The variation among the triplicate samples may be explained by the variable presence of atmospheric-derived oxygen and/or by varying microbial respiration, which will decrease the O_2_:Ar and increase δ^18^O_O2_ values. In fact, mixing and biological processes are likely both at work. Most remarkably, however, there is no plausible scenario to our knowledge whereby air contamination or microbial respiration could lead to the observed increasing trend in O_2_:Ar ratio coupled with decreasing δ^18^O_O2_ values. That is, air contamination would bring O_2_:Ar ratios and δ^18^O_O2_ back toward air-equilibrated values of ~23.88 ‰ (*49*) while microbial consumption of oxygen would lower O_2_:Ar while increasing δ^18^O_O2_ (*50*). The most parsimonious explanation for the observed trend would be the *in-situ* production of O_2_ with a very low δ^18^O (e.g., −20‰). Given the low δ^18^O (of H_2_O) of ground-waters in Alberta (−18.6 ‰ ± 2.0 ‰; mean ± SD; n=144, Data S1), which would be the source of oxygen for any *in situ* production of O_2_, even a small amount of production would have a large impact on lowering the overall δ^18^O_O2_ in groundwater, in the direction of our observations. In the absence of light, oxygen can be produced by water radiolysis (*51*), or microbially by chlorite dismutation (*52*), nitric oxide dismutation (*27*,*53*), and H_2_O_2_ dismutation (*54*).

**Fig. 8.**
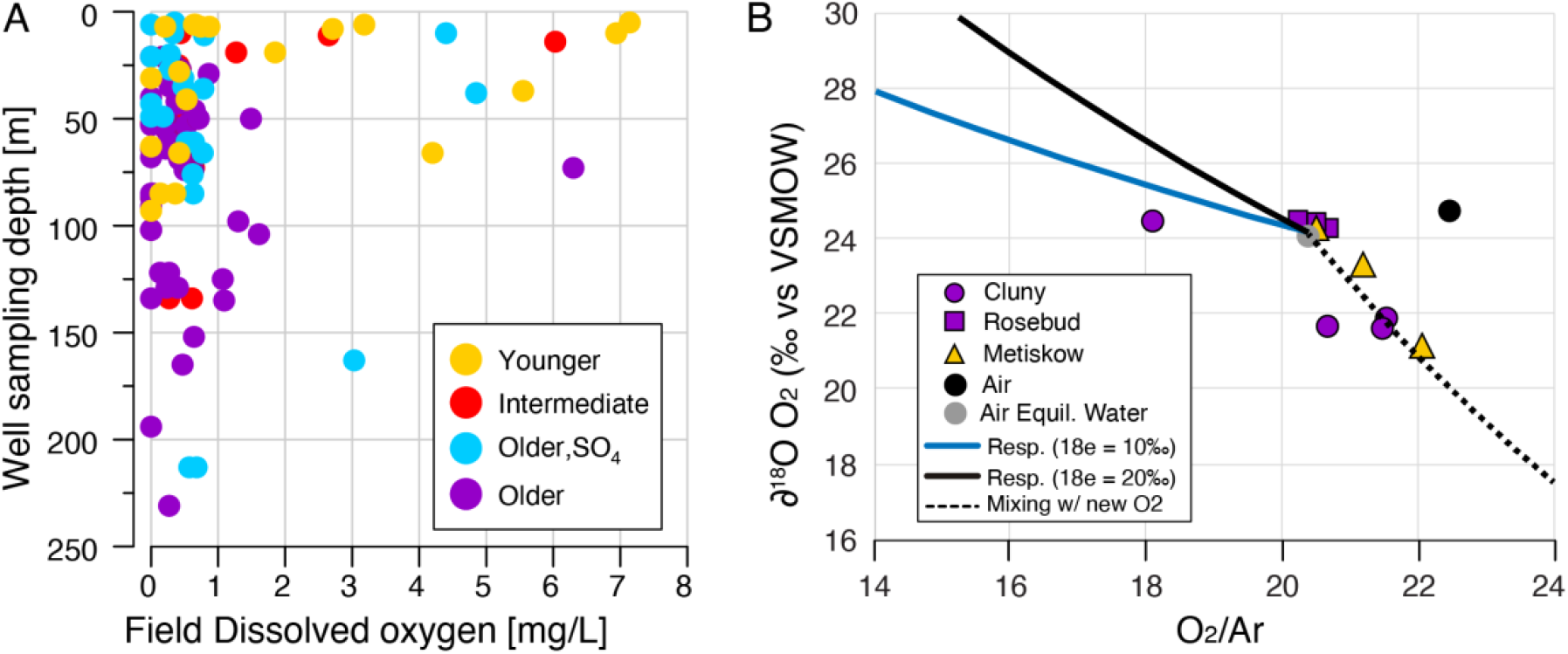
Concentration and isotopic signature of oxygen in groundwater samples. (**A**) Depth profile and dissolved oxygen concentration (mg/L) in the groundwater samples. (**B**) ^18^O-O_2_ isotopic signature over oxygen:argon ratio. The composition of lab-air equilibrated water and lab air are represented by a gray and black circle, respectively. Error bars are 1 standard deviation (n = 13). The two blue lines show different isotope effects that would be caused by fractionation due to respiration. During consumption of O_2_ by respiration (decreasing O_2_:Ar ratio), there is a preferential accumulation of ^18^O in the remaining O_2_ pool – leading to higher d^18^O values. As the degree of this isotope fractionation can vary the two solid lines represents two hypothetical, but realistic scenarios for how the O_2_ pool might change with microbial respiration. The black dashed line represents a mixing line between air equilibrated water mixed with a hypothetical ‘new source of O_2_’ that has a very low d^18^O (−20‰). Such light oxygen is consistent with the biological formation of O_2_.

## Microbial production of dark oxygen via dismutation

Microbes can produce O_2_ via chlorite dismutation, a process carried out among others by microbes of the genera *Dechloromonas* (*52*), *Dechlorobacter*, *Dechlorosoma*, *Azospira*, *Azospirillum* (*55*), *Nitrospina* and *Nitrobacter*, all of which were present in the GOWN aquifers based on the 16S rRNA gene analyses*. Dechloromonas* was among the most abundant genera of the study accounting for up to 30 % of all bacterial sequences in some samples. We performed metagenomic sequencing of samples from selected sites and reconstructed moderately abundant chlorite dismutase genes affiliating with *Dechloromonas, Nitrospina*, and *Nakamurella* species from old, hypoxic (0.5-0.63 mg/L) groundwaters of GW114 (Data S5). Groundwater can also contain a high diversity of nitric oxide dismutation genes (*56*) and indeed *Methylomirabilaceae* that dismutate nitrite were present in many samples. We have reconstructed a highly abundant nitric oxide dismutase gene (*nod*, at 700× coverage) from the metagenome of old, anoxic groundwaters of GW144 (Data S5). We found the *nod* gene in the assembled genome (98.5 % completeness, 0.5 % contamination) of an abundant *Hydrotalea* population (21% relative abundance). It was recently shown that nitric oxide dismutating *Nitrosopumilus sp*. can leak intracellularly produced oxygen into the medium possibly supporting other aerobic organisms (*53*). It was also reported that even *Pseudomonas aeruginosa* can dismutate nitric oxide and release peaks of up to 20 μM oxygen into their surroundings (*57*). Lineages affiliating with both, *Nitrosopumilus* and *Pseudomonas* were wide-spread and abundant in the studied groundwaters (*57*). Abundant membrane-bound dismutases in the studied aquifers and their release of oxygen into the surroundings could explain the observed presence of ^18^O-depleted oxygen in old groundwaters.

## Versatile microbial lineages, metabolisms, and niches in groundwater ecosystems

Our findings suggest that hydrogen fuels a rich mosaic of microbial metabolisms leading to extensive microbial productivity in the studied aquifer ecosystems (Fig 9), underlining the importance of hydrogen in the terrestrial subsurface (*58*). Key guilds in these ecosystems were hydrogenotrophic methanogens which provide methane for abundant methanotrophs. Substantial aerobic and anaerobic methane oxidation may reduce greenhouse gas emissions from coal beds and shales with relevance for carbon budgets and climate change. Extensive sulfur oxidation by thiotrophic microbes may reduce sulfide levels and H_2_S contents, which are produced by sulfate reducers in these aquifers. Microbial ecology and aqueous and isotope geochemistry suggest that groundwater microbial communities utilize the entire breadth of known biochemically available redox potential, from hydrogen to oxygen. Unlike for many other aquifers in which cell numbers decrease with groundwater age, cell numbers increased with age in groundwater obtained from aquifers in Alberta, corroborating that these aquifer communities are fueled by autochthonous energy sources derived from within the subsurface. Considering the size of the investigated area we conclude that global subsurface biomass may be underestimated, particularly in aquifers of organic carbon-rich strata, such as coal beds and shales. The geochemical and microbiological data suggest that oxygen is an important electron acceptor for microbial metabolisms in the studied aquifers. The presence of aerobic microbes in deep aquifers and bedrock has been previously reported (*59*), yet the mechanisms of deep oxygen migration or in situ production remain unclear. While co-migration of some oxygen with the aging groundwater could theoretically explain the observed microbial productivity, the presented oxygen isotope analyses suggest that at least some portion of the oxygen is generated in situ. Most of the hypoxic waters surveyed here are several thousand to more than 10,000 years old based on tritium and ^14^C data (Fig 2B), supporting that even deep subsurface ecosystems provide niches for aerobic microorganisms (*60*). The production of dark oxygen provides a mechanism for previously reported groundwater oxygen anomalies for which a compelling explanation was lacking thus far (*46*, *48*, *61*–*63*). Microbial dark oxygen production may be relevant for the evolution of the geobiosphere, as it provides a source of oxygen independent of light, on Earth as well as on other celestial bodies.

**Fig. 9.**
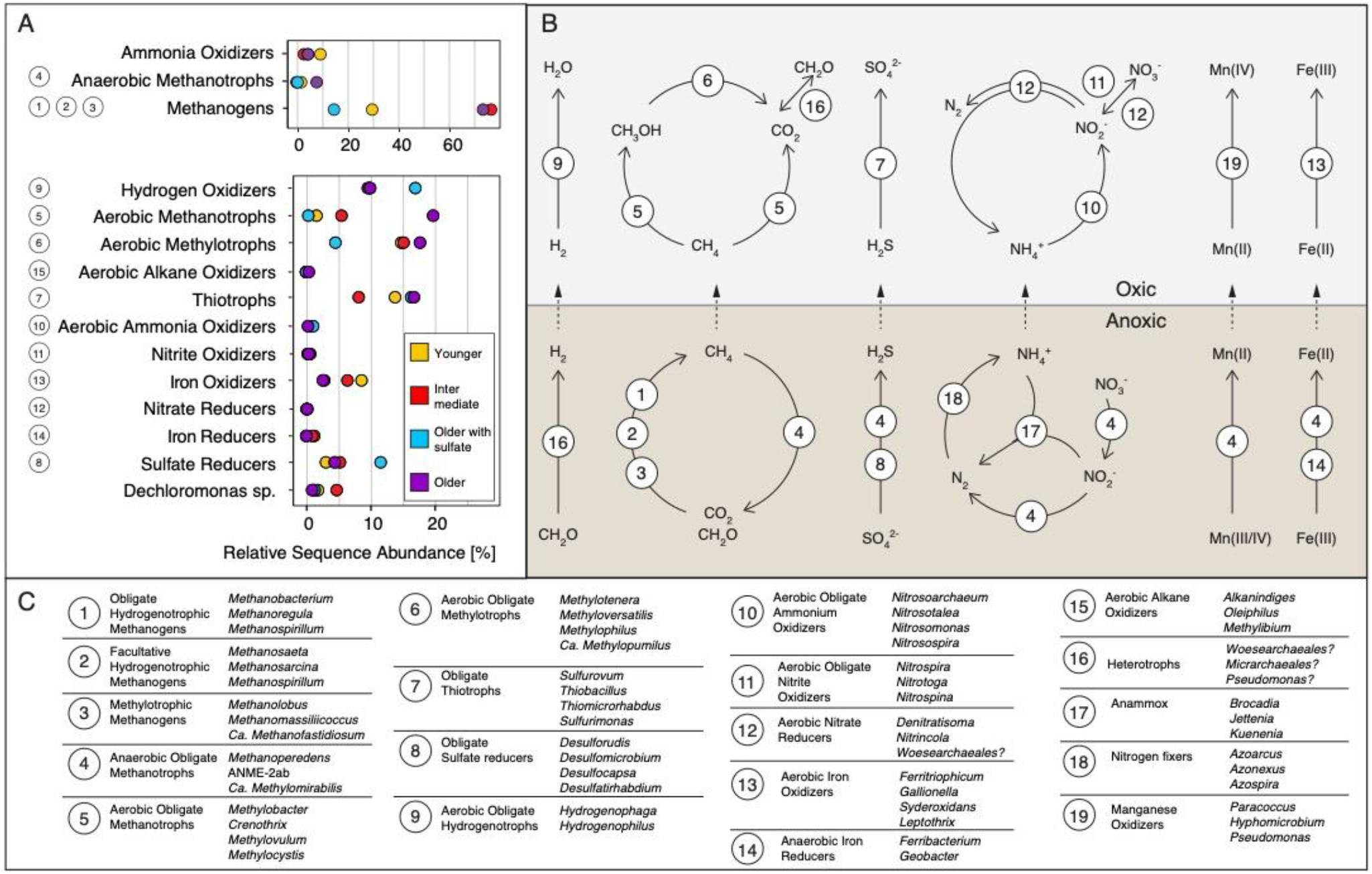
Microbial guilds and potential community functions. (**A**) Relative sequence abundances of microbial guilds. (**B**) Schematic overview of element cycling inferred from microbiological and geochemical analyses. Numbers refer to key microbial genera (**C**) with the potential to carry out the respective function. The list of microbes is not exhaustive and mainly contains the most abundant genera.

## Materials and Methods

### Groundwater wells

Approximately 40-90 groundwater wells are sampled annually for water quality by Alberta Environment and Parks (*8*) and analyzed for a variety of geochemical, isotopic, and microbial parameters. For this study we analyzed 159 groundwater samples from 95 monitoring wells, sampled from January 2016 to December 2017. 38 wells are screened within Neogene-Quaternary surficial deposits while 57 wells reach sedimentary bedrock, usually of Paleogene or Cretaceous age (Table S1). Well depths in this study ranged from five to 231 m (mean=60 m). The sedimentary bedrocks in which groundwater monitoring wells are completed include the Paskapoo and equivalent (n=25), Horseshoe Canyon and equivalent (n=13), Belly River Group (n=9), Bearpaw (n=4), Milk River (n=3) and Loon River (n=2) formations. One well was not assigned with a geological formation. For further details concerning the location and geology refer to the SI.

### Sampling

All wells were sampled at least once, while select wells were sampled during different seasons (up to three), or repeatedly to obtain sampling replicates (up to three). Some wells were installed as clusters, with multiple wells (up to three) at the same location completed at different depths within the same formation or across multiple formations. All wells were purged to improve sample quality (*64*) and samples were collected after field parameters (dissolved oxygen - DO, oxidation reduction potential - ORP, temperature, electrical conductivity - EC) had stabilized, indicating representative aquifer water. All wells were sampled following the same procedure and the samples were stored at 4 °C until further processing. Dependent on the location of the wells, the samples were in transit at 4 °C between 1-7 days.

Tritium (^3^H), and carbon-14 of dissolved inorganic carbon (^14^C_DIC_) samples were collected with a submersible pump (SP). Samples for other water quality parameters, isotopes, dissolved gas, and microbiological samples were collected with a bladder pump (BP). The SP is placed 2 m below the expected drawdown level for purging and sample collection. The submersible pump has polyethylene tubing that does not get cleaned, and Nalgene tubing that has been cleaned with a general laboratory detergent (Sparclean) followed by a rinse with ultrapure (18 MΩ) water. The BP is either placed in the vicinity of the SP or just above the screened interval for sample collection and has acid-washed, methanol and ultrapure water rinsed Teflon tubing. For microbiological analyses we collected 1000 ml groundwater in a sterile Nalgene bottle. The sample was sent to the laboratory at 4 °C and used for microbial cell cryo-preservation, microbial cell counts, and nucleic acid extractions.

### Chemical analyses

Groundwater samples for major cations and anions (including nitrate) were collected in polyethylene bottles and analyzed at ALS Global by Inductively Coupled Plasma-Mass Spectrometry (ICP-MS) and Ion Chromatography, respectively. Total and phenolphthalein alkalinity were determined by titration, and bicarbonate and carbonate concentrations were subsequently calculated. Dissolved organic carbon was measured by high temperature combustion on a total organic carbon analyzer with infrared detection. Dissolved trace element samples were filtered in the field using 0.45 μm in-line capsule filters, preserved to pH <2 with nitric acid, and analyzed at InnoTech Alberta by ICP-MS.

### Gas analyses

Dissolved gas samples were collected with a bladder pump. Evacuated crimp-top glass serum bottles with butyl septa and preserved with mercuric chloride were filled by piercing the septa with a needle. Gas composition was determined in the Applied Geochemistry Laboratory at the University of Calgary by gas chromatography. The gas dryness parameter is defined as the ratio between methane and higher n-alkanes. The isotopic composition of methane and CO_2_ was analyzed in the Isotope Science Laboratory at the University of Calgary on a ThermoFischer MAT 253 isotope ratio mass spectrometer (IRMS) coupled to a Trace GC Ultra and GC Isolink (ThermoFisher) and reported relative to V-PDB for δ^13^C and V-SMOW for δ^2^H. The precision for carbon isotope analyses was better than ± 0.5‰ for hydrocarbons, ±0.2‰ for carbon dioxide and better than ±2‰ for δ^2^H of methane.

### Groundwater dating

All samples for ^14^C_DIC_ analyses were shipped to the A.E. Lalonde Laboratory in Ottawa for analysis by accelerator mass spectrometry (AMS). Sample pretreatment techniques can be found in (*65*). Radiocarbon analyses are performed on a 3MV tandem accelerator mass spectrometer built by High Voltage Engineering (HVE). We used the fractionation-corrected fraction modern (the F^14^C value) for post-bomb ^14^C data according to the amended conventions (*66*). Tritium (^3^H) is a radioactive isotope of hydrogen with a half-life of 12.4 years. Tritium concentrations are measured in tritium units (TU) where 1 TU is defined as the presence of one tritium in 10^18^ atoms of hydrogen. For samples from the 2016-2017 sampling campaign, tritium in the obtained groundwater samples was analyzed at the A.E. Lalonde Laboratory in Ottawa by electrolytic enrichment and the standard method of liquid scintillation counting with a precision of 0.8 TU.

### Carbon, oxygen, nitrogen, and sulfur isotope analyses

Most stable isotope analyses were performed in the Isotope Science Laboratory at University of Calgary (https://www.ucalgary.ca/labs/isotope-science-lab/techniques). Dissolved inorganic carbon (DIC) isotope samples were filtered in the field using an in-line 0.2 μm capsule filter and collected in a 125 mL clear glass flint bottle with a cone cap. CO_2_ was produced from DIC by acid attack and δ^13^C was analyzed as described in the gas analyses section. For the isotopic composition of sulfate, samples were filtered in the field with an in-line 0.2 μm capsule filter and collected in 1 L wide mouth HDPE plastic bottles. Dissolved sulfate was converted into barium sulfate and analyzed using a ThermoQuest Finnigan Delta Plus XL IRMS coupled with a Fisons NA 1500 Elemental analyzer for δ^34^S_SO4_ and a HEKAtech HT Oxygen Analyzer with a Zero Blank autosampler for δ^18^O_SO4_. δ^34^S_SO4_ is reported relative to V-CDT and δ^18^O_SO4_ is reported relative to V-SMOW. Precisions for both δ^18^O_SO4_ and δ^34^S_SO4_ was ±0.5 ‰. The isotopic composition of nitrate was determined on N_2_O generated by the denitrifier technique using a Thermo Scientific Delta V Plus IRMS coupled with a Finnigan MAT PreCon. Precisions for δ^15^N_NO3_ and δ^18^O_SO4_ were ±0.3 ‰ and ±0.7 ‰ respectively.

### Oxygen isotope analyses on dissolved oxygen

Samples for the analysis of oxygen isotope ratios of dissolved O_2_ and O_2_/Ar ratio measurements in groundwater from monitoring wells were filled directly into 20 ml headspace vials allowing many volumes of overflow to displace air, poisoned with 100 μl of a saturated zinc chloride solution, quickly crimp-sealed with butyl septa (with no headspace bubble). At the lab, a 5 ml headspace was established by replacement with ultra-high purity helium. Vials were equilibrated with headspace on a shaker table at room temperature for several days. Aliquots of the headspace (containing O_2_ and Ar from the water sample) were directly injected into the sample inlet system (including a 2m, 5Å molecular sieve GC column) and routed into the isotope ratio mass spectrometer (Isoprime 100, multi-collector). Injections of lab air and lab-air equilibrated water were used as working standards. Based on repeat analyses the precision of the δ^18^O measurements was +/− 0.1‰ and of the O_2_/Ar was +/− 0.05.

### Cell staining, enumeration, and biomass yield calculations

To fix cells we added 4 ml formaldehyde solution (37 %) to 50 ml groundwater sample (f. c. ~2.7 %). The sample was stored at 4 °C until further processing. 1 ml sample was diluted in 10 ml 1×phosphate buffered saline (PBS) and filtered onto a polycarbonate filter (0.1 μm pore size, 25 mm diameter, Millipore Sigma) using a mixed cellulose ester membrane filter (0.45 μm pore size) as support. The filter was rinsed with 10 ml of 1×PBS, dried and stored until further processing. A section of each filter was stained with 1 μg ml^-1^ DAPI (4’,6’-diamidino-2-phenylindole) for 10 min at room temperature, washed with deionized water and 80 % ethanol and dried. Filter pieces were mounted on microscope slides using 4:1 Citifluor:Vectashield solution (VWR and Vector Laboratories) and stored at −20 °C. Cells were visualized and counted using an Axio Imager A2 (Zeiss, Jena, Germany) equipped with an X-Cite 120 LED (Excelitas, Waltham, USA) fluorescence light source and 12.5×12.5 mm ocular grid. 20-40 grids (at least 1000 cells) were counted per filter to obtain a robust dataset. To estimate the cell number than can be sustained per mol oxygen (Data S1) we used published values for microbial cell content (*67*) and biomass yield (*68*).

### 16S rRNA gene library preparation and ampli-con sequencing

Two to eight 50 ml groundwater sample tubes per sample were centrifuged at 4000 *g* for 1 hour at 4 °C. The supernatant was discarded, the pellets combined in a 2 ml tube and stored at - 80 °C until further processing. Genomic DNA was extracted using the DNeasy PowerLyzer PowerSoil Kit (12855-100, QIAGEN) according to manufacturer’s protocol with a minor modification; cells were lysed by bead beating at 4 m s^-1^ for 45 s using a Bead Ruptor 24 (OMNI). Extraction blanks were processed to detect potential laboratory contamination during extraction. DNA concentrations were measured fluorometrically using a Qubit 2.0 (Thermo Fisher Scientific, Canada). The bacterial 16S rRNA gene v3-4 region was amplified using S-D-Bact-0341-1-S-17 (5’-CCTACGGGAGGCAGCAG-3’) and a modified version of S-D-Bact-0785-a-A-19 (5’-GACTACHVGGGTATCTAATCC-3‘). The archaeal 16S rRNA gene v6-9 region was amplified using the primer pair S-D-Arch-0915-a-S-20/S-*-Univ-1392-a-A-15 (5’-AGGAATTGGCGGGGGAGCAC-3’, 5’-ACGGGCGGTGTGTRC-3’ (*69*). PCRs consisted of 8 μl (1-10 ng) DNA template, 2.5 μl of each primer (f.c. 1 μM), 12.5 μl 2× Kapa HiFi HotStart Ready Mix (Kapa Biosystems, Wilmington, MA, USA) and PCR-grade water ad 25 μl. For bacteria, a touchdown PCR program was used for improved annealing: initial denaturation at 95 °C for 3 min, 10 cycles of 95 °C for 30 sec, 60 °C for 45 sec (touchdown −1 °C per cycle), 72 °C for 60 sec, followed by 20 cycles of 95 °C for 30 sec, 55 °C for 45 sec, 72 °C for 60 sec, and a final extension at 72 °C for 5 min. For archaea, the touchdown PCR program was: initial denaturation at 95 °C for 5 min, 10 cycles of 95 °C for 30 sec, 62 °C for 45 sec (touchdown −1 °C per cycle), 72 °C for 60 sec, followed by 20 cycles of 95 °C for 30 sec, 60 °C for 45 sec, 72 °C for 60 sec, and a final extension at 72 °C for 5 min. PCRs were performed in triplicate, pooled, and purified using 56 μl Agencourt AMPure XP beads (Beckman Coulter, Indianapolis, USA) per pooled PCR product (~65-75 μl) following manufacturer’s instructions. Amplicons were indexed using 5 μl purified PCR product, 5 μM of each Index Primer (f.c. 1 μM), 25 μl 2× Kapa HiFi HotStart Ready Mix and 10 μl PCR-grade water. Indexing PCR program: initial denaturation at 95 °C for 3 min, 10 cycles of 95 °C for 30 sec, 55 °C for 45 sec, 72 °C for 60 sec, and a final extension at 72 °C for 5 min.

Indexed amplicons were purified with Agencourt AMPure XP beads. The concentration and size of indexed amplicons were checked with a Qubit 2.0 fluorometer and Agilent 2100 Bioanalyzer system (Agilent Technologies, Mississauga, ON, Canada), respectively. Indexed amplicons were pooled in equimolar amounts and sequenced using Illumina’s v3 600-cycle (paired-end) reagent kit on an Illumina MiSeq benchtop sequencer (Illumina Inc., San Diego, CA, USA) after all DNA extraction blanks and PCR reagent blanks were confirmed for negative amplification.

### Community analyses

Raw sequences were analyzed using *DADA2* v1.16 (*70*). Briefly, forward, and reverse reads were quality-trimmed to 275 bp and 205 bp, respectively, and primer sequences (17 bp forward, 21 bp reverse) were removed. Reads with more than two expected errors were discarded, paired reads were merged, and chimeric sequences were removed. Species level taxonomy was assigned with silva_nr_v138_train_set and silva_species_assignment_v138. After quality control and the removal of blanks, biological and technical replicates we obtained 64 archaeal and 110 bacterial amplicon datasets. Archaeal datasets contained a total of 3.89 × 10^5^ sequence reads belonging to 2633 unique amplicon sequence variants (ASVs). Archaeal samples had on average 6083 ± 6387 reads (mean ± standard deviation) and 69 ± 81 ASVs (Table S2). Bacterial datasets comprised a total of 4.65 × 10^6^ sequence reads belonging to 14665 unique ASVs. Bacterial samples had on average 4.23 ± 1.57 × 10^4^ reads and 272 ± 221 unique ASVs (Table S3). Archaeal and bacterial ASV nucleotide sequences and taxonomy are listed in Data S6 and S7, respectively. The archaeal or bacterial ASV-by-sample table (Data S3, S4) was used to determine the number of observed ASV, absolute singletons, relative singletons, relative abundance, and composition. Alpha diversity (richness, Shannon entropy, Inverse Simpson Diversity and Chao1 estimated richness) was calculated from the ASV-by-sample table using a subsampling approach to account for unequal sampling effort. We used 1008 and 6796 randomly chosen reads from each archaeal and bacterial sample, respectively. In the case of archaea about half of the samples had less reads and were thus removed from the analysis. Differences in diversity between conditions were tested using the Wilcoxon signed rank-test (*ggsignif*) as implemented in *ggplot2* (*71*). Bray-Curtis dissimilarities between all samples were calculated and used for two-dimensional non-metric multidimensional scaling (NMDS) ordinations with 20 random starts. Environmental parameters that significantly impacted the archaeal (Table S4) and bacterial community (Table S5) based on *envfit* were chosen for a Redundancy Analyses using Hellinger-transformed ASV data. Analyses were done with VisuaR v02 (https://github.com/EmilRuff/VisuaR) a workflow based on the R statistical environment, including the packages *vegan* (*72*), *ggplot2* (*71*) as well as custom R scripts. To support the reliability of the results we determined that storage duration (Fig. S12), DNA extraction, and sample handling did not affect the community structure (not shown).

### Metagenome sequencing and analysis

Metagenomes were processed and sequenced at the Center for Health Genomics and Informatics in the Cumming School of Medicine, University of Calgary. Genomic DNA was sheared into ~350 bp fragments using a S2 focused ultra-sonicator (Covaris, Woburn, MA). Libraries were prepared using the NEBNext Ultra II DNA Library Prep Kit for Illumina (New England Biolabs, Ipswich, MA) according to the manufacturer’s protocol, including size selection with SPRIselect magnetic beads (Beckman Coulter, Indianapolis, IN) and PCR enrichment (eight cycles) with NEBNext Multiplex Oligos for Illumina (New England Biolabs, Ipswich, MA). DNA concentrations were estimated using qPCR and the Kapa Library Quantitation Assay for Illumina (Kapa Biosystems, Wilmington, MA). Genomic DNA was sequenced on an Illumina NovaSeq 600 sequencer (Illumina, San Diego, CA) using a 300 cycle (2 × 150 bp) S1 flow cell. Quality control was performed on raw, paired-end Illumina reads using BBDuk, including trimming, filtration of contaminants, and clipping low-quality ends. Reads that passed quality control were assembled into contigs using Megahit (*73*) and contigs of < 500 bp were not included in subsequent steps. Reads from each sample were mapped to each of the 25 samples using BBMap and depth profiles were generated using the ‘jgi_summarize_bam_contig_depths’ function from MetaBat2 (*74*). Assembled contigs were binned into metagenome-assembled-genomes (MAGs) using MetaBat2, CONCOCT (*75*) and MaxBin2 (*76*). DAS-Tool (*77*) was used to integrate MAGs produced by the three binning tools. The contamination and completeness of MAGs were assessed by CheckM v1.2.0 (*78*). MAG classification using GTDB-tk (version 2.1.0, database release r207) (*79*). Transfer RNA, ribosomal RNA, CRISPR elements, and protein-coding genes including nitric oxide dismutase coding genes and perchlorate dismutase coding genes were predicted and annotated using MetaErg (*80*).

## Data and materials availability

Archaeal and bacterial 16S rRNA amplicon data are publicly available under SRA BioProject PRJNA861683. Shotgun metagenomic data and metagenome-assembled genomes (MAGs) are available under BioProject PRJNA700657. Comprehensive contextual data are publicly available from the publisher for Earth and environmental data PANGAEA (accession number pending).

## Acknowledgements

We are very grateful to Joanna Borecki, James Rogans, Dennis Rollag, Vien Lam and other staff of the Groundwater Observation Well Network of Alberta Parks and Environment (https://www.alberta.ca/groundwater-observation-well-network.aspx) for providing access to groundwater monitoring wells, for sampling and for sharing measurement results and expertise. Without their ability to obtain highest quality groundwater samples and their input this study would not have been possible. We also greatly appreciate the support and expertise of Carmen Li regarding nucleic acid sequencing. We thank Manuel Kleiner and Xiaoli Dong for support and discussions concerning all things bioinformatics. We thank Anirban Chakraborty for insights on sample handling, Steven Taylor and Veith Becker for support with isotopic measurements and interpretation, and Rachel HR Stanley for help with isotope mass spectrometry.

## Funding

Alberta Innovates Technology Futures (AITF)/Eyes High Postdoctoral Fellowship (SER)

Start-up funds by the Marine Biological Laboratory, Woods Hole (SER)

Alberta Innovates Energy and Environment Solution (AIEES) - Project “geochemical resource characterization of Alberta groundwater” (BM)

Alberta Innovates Water Innovation Program (AI-WIP) - Project “occurrence, origin and fate of aqueous contaminants in Alberta groundwater” (BM)

## Author contributions

Conceptualization: SER, MS, BM

Methodology: SER, MS, PH, CNM, IHdA, SDW, AS

Investigation: SER, PH, IHdA, MN, MD, SC, LC, OOK, AS, SB, SDW

Formal Analysis: SER, PH, IHdA, CNM, MD, SDW

Visualization: SER, PH, IHdA, SDW

Supervision: SER, PH, BM, MS

Writing—original draft: SER, PH

Writing—review & editing: All coauthors

## Supplementary Materials

### This bioRxiv preprint PDF file includes

Supplementary Text

Figs. S1 to S12

Tables S1 to S5

References (81 to 107) (if applicable—these should refer only to references in the SM)

### Other Supplementary Materials for this manuscript will be available at PANGAEA soon (submission in progress) and include the following

Data S1. Physical, chemical, isotopic, and microbiological data of the studied GOWN wells

Data S2. Fluorescence micrographs of groundwater communities

Data S3. Sequence abundance of archaeal 16S rRNA gene amplicon sequence variants

Data S4. Sequence abundance of bacterial 16S rRNA gene amplicon sequence variants

Data S5. Amino acid sequences of dismutase genes

Data S6. Nucleotide sequences and taxonomy of archaeal amplicon sequence variants

Data S7. Nucleotide sequences and taxonomy of bacterial amplicon sequence variants

## Supplementary Text

### Geologic formations

Situated in the Western Canadian Sedimentary Basin, Alberta’s conventional and unconventional hydrocarbon reserves include bitumen in northern Alberta (i.e. Athabasca oil sands deposits in Fort McMurray, Cold Lake and Peace River), shale gas and tight gas in the northwest (i.e. Duvernay shale, Montney tight sandstone/shale, Muskwa formation), and coal bed methane (CBM) in south-central Alberta (i.e. coal zones from the Horseshoe Canyon Formation, Belly Group, Mannville Group) (*11*) (Fig. 1). The terrestrial fluvial Paskapoo Formation consists of mudstone and siltstone, with thick tabular sandstones deposited in fluvial channels (*13*, *81*, *82*). The terrestrial fluvial Horseshoe Canyon Formation consists primarily of sandstone, siltstone, and coal. The marine Bearpaw formation interfingers with the lower and middle Horseshoe Canyon formation and consists predominantly of shale and minor sandstone and coal (*81*). The Belly River Group consists of fluvial sandstone and siltstone with minor mudstone and coal. The coal zones with coal bed methane (CBM) potential are included in the Horseshoe Canyon and the Belly River formations. Thin coal seams may also occur in the Paskapoo Formation. All these bedrock formations comprise heavily used aquifers in the prairies region of Alberta (*83*) although they are highly heterogeneous. Neogene-Quaternary surficial deposits of varying thickness and composition (e.g., till) overlie bedrock and are associated with glacial retreat. Glacial sand and gravel deposits that fill buried river valleys (channels) are also commonly used aquifers in Alberta (Table S1). Consequently, the GOWN wells (Groundwater Observation Well Network) are completed in many different lithologies including sandstone, siltstone with intermittent coal or shale beds, pre-glacial sand, and sandy and gravelly lacustrine or moraine deposits. More information about these geological formations can be found in (*13*, *81*–*87*).

### Hydrochemical facies

Hydrochemical facies are defined as groundwater masses that have different geochemical attributes (*88*). These hydrochemical facies are frequently compared using graphical representation (*89*). The traditional Piper diagram in Fig. 2A indicates that there are different hydrochemical facies with the groundwater samples varying from Ca-Mg-HCO_3_ to Na-HCO_3_, SO_4_-rich water as well as mixing water facies. Access to groundwater data using radioisotopes such as tritium and ^14^C_DIC_ permits to estimate the travel time between the point of recharge and the point of sampling (*12*). The chemical evolution using the Ca/Na mass ratios proxy and preliminary age dating data permit to delineate further those hydrochemical facies (Fig. 2B). In this study, the youngest tritium-containing groundwater samples (indicating recharge after 1960) were characterized by low total dissolved solids (< 400 mg/L), high Ca/Na ratio (median of 3.5) and a calcium-magnesium-bicarbonate dominated water chemistry resulting from dissolution of carbonates during recharge (yellow symbols, Fig. 2). Those groundwater samples were collected from wells completed mainly in Neogene-Quaternary surficial deposits. In contrast, the most evolved groundwater samples containing no tritium and having ^14^C_DIC_ ages (uncorrected) indicative of groundwater more than several hundreds of years old, had elevated average contents of total dissolved solids (>900 mg/L) and a low Ca/Na ratio (median of 0.01). Old waters were rich in sodium, bicarbonate, and chloride due to water-rock interactions including ion exchange, and weathering of minerals (purple symbols, Fig. 2). Those groundwater samples were collected from wells completed mainly in buried valleys, Paskapoo, Horseshoe Canyon and Milk River formations. Sulfate derived from oxidation of reduced-S, anhydrite or gypsum dissolution is ubiquitous in a third group of groundwater samples characterized by variable groundwater ages, elevated total dissolved solids (> 1700 mg/L) and low Ca/Na (median of 0.12) (blue symbols, Fig. 2). These sulfate-rich waters are mainly associated with sodium-sulfate, sodium-bicarbonate-sulfate hydrochemical facies (low Ca/Na). A minority of samples is associated with calcium-sodium-bicarbonate-sulfate hydrochemical facies resulting in higher Ca/Na ratios. Those groundwater samples were collected from wells completed in Neogene-Quaternary surficial deposits or in the Cretaceous Paskapoo, Horseshoes Canyon, Bearpaw, Loon River formations. The groundwater mixture often results in intermediate hydrochemical facies (red symbols, Fig. 2) associated with calcium-sodium-bicarbonate-chloride waters. These groundwater samples are collected from wells mainly completed in Neogene-Quaternary surficial deposits.

### ^14^C-based Groundwater dating

Reimer et al. (*66*) point out the problems with the different conventions in radiocarbon measurements for post-bomb ^14^C data and define a fractionation-corrected fraction modern (the F^14^C value) according to the amended conventions (*90*):

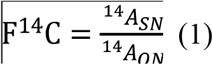

where ^14^A_SN_ and ^14^A_ON_ refer to the fractionation-corrected or normalized ^14^C activity for the sample and the secondary Oxalic acid-II (Ox2) standard used for the ^14^C-measurements (*65*) respectively. Both specific activities of the sample and Ox2 ^14^A_S_ and ^14^A_ox2_ are first measured and then normalized to ∂^13^C= −25‰ giving the normalized sample activities ^14^A_SN_ and ^14^A_ON_ as follow:

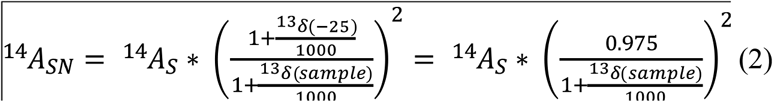

For the standard according to Mook and van der Plicht (*91*)

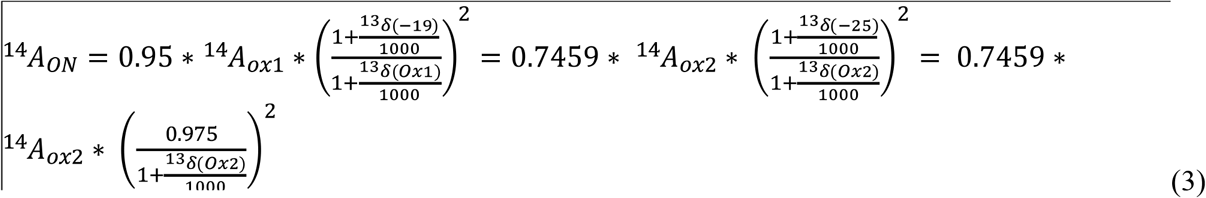

Thus, from equations (1), (2) and (3):

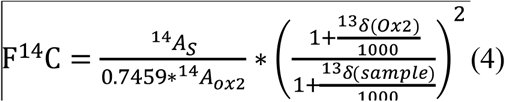

F^14^C can thus ranges between 0 and 1 (1 for modern value) or 0 to 100 % modern carbon.

The conventional radiocarbon age is given by:

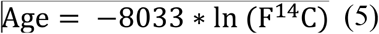

with the half-life value T_1/2_ of 5568 years divided by ln(2) resulting in the value 8033. The ^14^C ages thus calculated are reported in BP or “Before Present” the term “present” does not correspond to the present day but it refers to the standard activity 1950 AD. The measured activity plus error F^14^C ±σ(F^14^C) reported can be translated into an age T± σ(T).

#### Younger, recently recharged groundwater

A total of 17 % (20 out of 112) groundwater samples fell in the Ca^2+^-Mg^2+^-HCO_3_^-^ (calcium-magnesium-bicarbonate) hydrogeochemical facies (yellow symbols; Fig. 2A). The water has on average a low concentration of Total Dissolved Solids (TDS) of 361 ± 125 mg/L (n=20). Among the alkaline earths, the concentration of Ca^2+^ and Mg^2+^ ions ranged from 26.8 to 142 mg/L and 13.0 to 52.9 mg/L with a mean of 77.8 ± 29.9 mg/L and 27.1 ± 10.2 mg/L (n=20) respectively. Sodium concentrations were on average 24.7 ± 8.8 mg/L. Such facies are marked by a mass ratio Ca^2+^/Na^+^ > 0.7 (Fig. 2B). For the anions, the average concentrations of HCO_3_^-^, SO_4_^2-^, NO_3_^-^, Cl^-^ are 34.5 ± 77.3 mg/L, 37.5 ± 42.2 mg/L, 10.6 ± 26.7 mg/L, 10.5 ± 21.9 mg/L (n=20) respectively. Values of dissolved inorganic carbon δ^13^C_DIC_ ranged from −12 ‰ to −16 ‰ suggesting carbonate minerals dissolution in closed and open systems, respectively. Environmental tracers such as ^14^C, ^3^H have helped refine estimates of recharge, flow time scales and “groundwater age”. “Groundwater age” refers to the travel time between the point of recharge and the point of sampling. A subset of nine samples has been submitted for ^3^H and ^14^C_DIC_ analyses. The relatively high ^3^H content (mean of 9.5 Tritium Units (T.U.), n=7) indicates that these waters had a relatively short residence time (post-year 1953). Only two out of nine younger waters had ^3^H values below detection limit <0.8 T.U. F^14^C_DIC_ had an average of 68.2 ± 15.3 (n=13) with fractions varying from 40 to 84 % modern carbon, similar to previously characterized young groundwater of 85±5 % modern carbon (*12*).

#### Groundwater of intermediate age

A total of 10 % (11 of 112) groundwater samples fell into mixed groundwater hydrochemical facies dominated by Na^+^, Ca^2+^ and/or Mg^2+^ and HCO_3_^-^ and/or Cl^-^ (red symbol, Fig. 2A). The water had an average TDS concentration of 572 ± 243 mg/L (n=11). The concentrations of Na^+^, Ca^2+^ and Mg^2+^ were on average 87.2 ± 27.9 mg/L, 72.8 ± 31.5 mg/L, 44.1 ± 36.1 mg/L respectively, with mass ratios of 0.3 < Ca^2+^/Na^+^ <1.1 (Fig. 2B). HCO_3_^-^ content was 483 ± 196 mg/L, SO_4_^2-^ was 66.3 ± 70.3 mg/L, Cl^-^ was 56.6 ± 120 mg/L, and NO_3_^-^ was only detected in 4 samples. When detected, the concentration of NO_3_^-^ was 0.13 mg/L. A subset of 7 groundwater samples out 12 were submitted for ^3^H and ^14^C_DIC_ analyses. While the F^14^C_DIC_ values show a wide range from <0.5 (detection limit) to 80 % modern carbon, few groundwater samples are associated with ^3^H > 0.8 TU (n =2, up to 11.7 TU).

#### Older, geochemically mature groundwater

A total of 51 % (57 of 112) groundwater samples fell into the Na^+^-HCO_3_^-^-Cl^-^ (sodium-bicarbonate-chloride) hydrochemical facies (purple symbols; Fig. 2A). The water had an average TDS concentration of 1103 ± 889 mg/L (n=57). Sodium was the dominant cation with an average concentration of 427 ± 359 mg/L. A significant decrease in the concentration of Ca^2+^ was observed with an average of 12.8±21.3 mg/L. The mass ratio Ca^2+^/Na^+^ was <0.5 (Fig. 2B). Bicarbonate concentration was 664 ± 343 mg/L (n=112), Cl^-^ was on average 67 ± 126 mg/L, with 7 outlier samples featuring concentrations of Cl^-^ > 500 mg/L. Sulfate was in average < 131 mg/L, nitrate was not detected except in two samples containing nitrate concentrations of > 3 mg/L. A subset of 30 groundwater samples was submitted to ^3^H and ^14^C_DIC_ analyses. The ^3^H content of 90 % of old groundwater samples was below the detection limit, only three samples showed a ^3^H > 0.8 TU. The ^14^C activity of 8 old groundwater samples was below the detection limit of the radiocarbon technique, while 22 samples had an average F^14^C_DIC_ of 17.8 ± 17.7 modern carbon.

##### Older, sulfate-rich groundwater

A total of 21 % (24 of 112) groundwater samples fell into hydrochemical facies dominated by SO_4_^2-^ (blue symbols; Fig. 2A). These waters had an average TDS concentration of 2882±3555 mg/L directly correlated with average SO_4_^2-^ concentrations of 1563±2423 mg/L. The average content of HCO_3_^-^ is 650 ± 287 mg/L. The major cation concentrations varied and allowed to identify two sub-categories of sulfate-rich waters, those dominated by Ca^2+^ and those dominated by Na^+^. In the former, the calcium concentration was 139 ± 79.2 mg/L (n=4), magnesium was 54.4 ± 30.1 and sodium was 92.2 ± 90.0 mg/L. Resulting in a mass ratio of Ca^2+^/Na^+^ > 1 (Fig. 2B). The hydrochemical type of these waters develops into Ca^2+^-Mg^2+^-HCO_3_^-^-SO_4_^2-^. In the Na^+^-rich waters, the sodium concentration was 870 ± 915 mg/L (n=20), calcium was 93.8 ± 130 mg/L (n=20), and magnesium was 85.8 ± 183. This results in a mass ratio of Ca^2+^/Na^+^ <0.8. The hydrochemical type of the water develops into Na^+^-(HCO_3_^-^)-SO_4_^2-^. A subset of 8 groundwater samples was submitted for ^3^H and ^14^C_DIC_ analyses. While the F^14^C_DIC_ values showed a wide range from <0.5 (detection limit) to 80 % modern carbon, few groundwater samples were associated with ^3^H > 0.8 TU (up to 8.3 TU). For both subtypes, the δ^13^C_DIC_ values had a narrower range between −20 ‰ to −12 ‰ indicating that most of the DIC content resulted from organic matter oxidation and/or carbonate mineral interactions in these systems.

### Methane and higher alkanes

Methane was present in all groundwater samples in a wide range of concentrations from 0.006 to >74 mg/L. The average concentration of methane were 6.87±15.9 mg/L and 0.03 mg/L respectively (n=106). Of the groundwater samples containing methane 73 % had a concentration of <1 mg/L. Ethane was detected in 23 samples with mean and median concentrations of 35 ± 71 μg/L and 2 μg/L respectively. Methane concentrations were significantly higher in old waters compared to other waters (p<0.05) (Fig. S6A-C) and were often associated with low concentrations of oxygen, nitrate, and sulfate (Fig 3A, 4, Table S1). Isotopic analysis supported widespread microbial methane production with δ^13^C_CH4_ < −60‰, δ^2^H_CH4_ < −200 ‰. Especially in older waters δ^13^C_DIC_ > +10‰ strongly supported methane production via CO_2_ reduction (Fig 3C), which is supported by the presence of high relative sequence abundances of hydrogenotrophic methanogens (Fig S4). Aceticlastic and methylotrophic methanogens were less abundant in older waters and largely occurred in younger and intermediate waters, as well as numerous sulfate-rich aquifers (Fig. S4–S7). A total of 17 samples with an average low methane concentration of 0.03 mg/L had increased δ^13^C_CH4_ values of over −50 ‰ up to −22 ‰ suggesting that methane oxidation has occurred. Methane oxidation apparently occurred in many groundwater samples that also showed sulfate reduction. In groundwaters with low concentrations of electron acceptors, the in-situ production of methane by methanogenesis becomes a thermodynamically favorable process. Methanogenesis is believed to proceed through two different major pathways, which are the reduction of inorganic carbon in the presence of hydrogen (hydrogenotrophic methanogenesis; Eq. (1)):

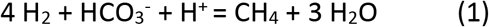

or the fermentation of acetate (acetoclastic methanogenesis; Eq. 2):

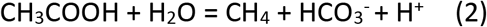

### Sulfate

Sulfate was widespread in Albertan groundwater samples and often present in relatively high concentrations as compared to the average standard sulfate concentration (250 mg/L WHO). Natural sources of sulfate include sulfur mineral dissolution, atmospheric deposition, and sulfide oxidation from minerals (*92*). Reduced sulfur in minerals such as pyrite or in organo-sulfur compounds are common in shales, coal seams and many other rock types. One hypothesis is that the oxidation of reduced sulfur (pyrite and/or organo-sulfur compounds), incorporated from the Albertan bedrock into the glacial deposit during glaciation, occurs in the till (*93*). Oxygen dissolved in the recharging groundwater and gaseous oxygen created oxic conditions needed to convert reduced sulfur into sulfate. S- O- isotope ratios of sulfate have been investigated to further evaluate the possible origin of sulfate in the groundwater. The δ^34^S_SO4_ values in groundwater ranged from −7.4 to +65.2 ‰ with a mean value of +1.6 ‰ (n=89). The δ^18^O_SO4_ values in groundwater ranged from −16.1 to +15.3 ‰ with a mean value of – 1.4 ‰ (n=89). The highest concentrations of sulfate were found in sulfate-rich, old groundwater that had δ^18^O_SO4_ values of below 0 ‰ and δ^34^S_SO4_ values of below 0 ‰ (Figure 3 A, B), suggesting that indeed oxidation of sulfide minerals was the main sulfate source. It is known that the light isotopes ^32^S and ^16^O are preferentially metabolized during bacterial sulfate reduction, thus causing a decrease in sulfate concentrations while ^34^S and ^18^O become progressively enriched in the remaining sulfate. We show that many samples with low sulfate concentrations had increased δ^34^S_SO4_ values > +10 ‰ suggesting that sulfate reduction has occurred in the aquifers (Fig. 3A). These waters also showed elevated δ^18^O_SO4_ values and often high methane concentrations.

### Oxygen

Highest dissolved oxygen (DO) concentrations were found in the freshly recharged (Ca^2+^-Mg^2+^-HCO_3_^-^) groundwater with an average concentration of 1.83 ± 2.35 mg/L, up to 7 mg/L approaching the saturation of oxygen in water, indicating mixed oxic-hypoxic conditions. Intermediate DO concentrations of 1.18 ± 1.76 mg/L were measured in intermediate, and sulfate-rich old groundwater samples (0.88 ± 1.29 mg/L), and lowest average DO of 0.51 ± 0.87 mg/L was associated with sulfate-poor, old water (Na^+^-HCO_3_^-^-Cl^-^). At certain sites we observed low δ^18^O_O2_ values (as low as +21 ‰), while also finding elevated O_2_:Ar ratios (Fig. 8B). The lower δ^18^O_O2_ and higher O_2_:Ar ratios interestingly fell along a trend, consistent with the simulated addition of DO with a δ^18^O value much lower than that of air-equilibrated water (dashed trend line in Fig 8B). As an idealized example, if 0.5 mg/L of O_2_ with δ^18^O_O2_ equal to −20‰ were added to air-equilibrated water, the net effect would be to lower the δ^18^O_O2_ by ~2‰ while increasing the O_2_:Ar ratio by ~5%, consistent with the slope of the dashed line in Fig 8B.

### Nitrogen compounds, iron, and manganese

In our study nitrate was only detected in younger groundwater, although it is not uncommon even in aquifers bearing older groundwater, pre 1953 (*94*). Nitrite was detected in 4 samples (<0.076 mg/L). The dissolved Fe and Mn concentrations in the groundwater samples are generally low, both mostly below 0.2 mg L^-1^ with maximum concentrations of 1.5 mg L^-1^ and 6.7 mg L^-1,^ respectively.

### Cell abundance

In addition to enumerating the cells in 78 samples (Fig 4) we have estimated the number of cells that can be sustained by the measured dissolved oxygen concentration in the water samples (Table S2). We used previously published values for biomass per mol oxygen yields (*68*) and biomass per microbial cell (*67*). The formula is included in the supplementary Table S2. We found that the counted cell number could have been sustained by the measured oxygen concentration in almost all aquifers, indicating that oxygen was not limiting in these systems.

### Beta diversity – Community Shifts

Any two samples shared on average 5.7 % of archaeal and 5.5 % of bacterial ASVs (Figure 7). There were, however, very different samples that shared no ASV and similar samples that shared 95 % archaeal or 71.4 % bacterial or ASVs. The archaeal and bacterial community structure of all four water types is different yet overlapping (ANOSIM: R_Arc_=0.29, p_Arc_=0.001, R_Bac_=0.33, pBac=0.001), while younger and older groundwaters are significantly different (R, p), indicating substantial community turnover with depth (Figure 7, S3). However, some microorganisms are exchanged between aquifers, because around 1 % of all archaeal (32 ASV) and 3 % of all bacterial ASV (388 ASV) occurred in all water types. The patterns of ASV occurrence also indicate that intermediate and old waters are in exchange with young waters (Figure S3), because the number of ASVs that exclusively occur in young and old waters (87 archaeal and 234 bacterial ASV) is higher than the number of ASV that exclusively occur in intermediate and old waters (10 archaeal and 157 bacterial ASV).

### Community variation

We investigated which environmental parameters are likely impacting the microbial community structure. We used an environmental fitting model (EnvFit in R package vegan) and calculated significance values (Table S7, S8) to select candidates for an in-depth Redundancy Analysis (RDA, Figure 5C, D). Both EnvFit and RDA, as well as partial RDA analyses rely on statistical significance testing using permutation tests (*95*). Groundwater microbial diversity was apparently influenced by so many factors, that even the most significant such as methane and carbonate could only explain a few percent of the total variation. Of the 30 physicochemical properties that we measured 14 significantly influenced archaeal populations, i.e., can be seen as factors that explain variation in archaeal community structure, while eleven were significant for bacteria (Table S7, S8, based on ASV data, EnvFit p<0.01). We took the eleven factors that were most important for both archaea and bacteria for further testing using RDA. The full model was highly significant for both domains and explained 24 % of archaeal and 18 % of bacterial variation. Upon investigating each parameter in a reduced model, we found that variation in archaeal community structure was explained by dissolved oxygen concentration and ^13^C-DIC at low significance (Fig 5A). Bacterial community structure was explained by methane concentration with very high significance, but well depth, hardness, calcium, and magnesium concentrations also played a significant role (Fig 5D).

### Community composition

#### Methanogens

The large majority of archaeal ASV affiliated with lineages known to perform methanogenesis (Figure 8A). Hydrogenotrophic methanogens were found in all water types and in all samples with few exceptions (GW128, GW3002, GW220, GW104, and GW259). Here, the methanogenic community consisted of methylotrophic clades, mainly *Methanolobus sp*. and *Methanomassiliicoccales*. Overall, hydrogenotrophic *Methanobacterium sp*. and *Methanoregula sp*., as well as aceticlastic *Methanosaeta sp*. were most abundant in the datasets. Although methylotrophic clades occurred at high relative abundances (> 50 %) in four samples of mature waters, they were generally more abundant in younger and sulfate-rich waters. Methanogenic archaea apparently tolerated low concentrations of oxygen that were present in most aquifers (Figure 4).

#### Methanotrophs, methylotrophs and short-chain alkane oxidizers

ASVs affiliating with *Methanoperedenaceae* – known to perform the anaerobic oxidation of methane (AOM) – almost exclusively occurred in old waters. In three aquifers (GW218, GW111 and GW456) these methanotrophs accounted for over 50 % of the archaeal community indicating that AOM may be of significance in these aquifers. ASVs affiliating with clades that are known to perform aerobic metabolisms accounted for most of bacterial diversity (Figure 8B). Aerobic methylotrophic *Methylophilaceae* occurred in all waters except those with sulfate, while *Methylomonadaceae* predominantly occurred in older waters. *Methylibium sp*. and obligate alkane-degrading *Alkanindiges sp* indicate that short-chain alkanes and aromatics could serve as a carbon and energy source in some of the aquifers (*96*, *97*). Despite the large overall diversity in methane-cycling clades each investigated aquifer contained signatures of at least one methanogen, methanotroph and methylotroph, supporting the potential for methane remediation and indicating that these aquifers are biofilters removing methane before it outgasses to the atmosphere. The presence of aerobic and anaerobic methanotrophs was reported for several deep bedrock sites (*59*). Their coexistence and the resulting syntrophy as well as functional redundancy suggest that methane cycling is an essential ecosystem function in high productivity aquifers.

#### Sulfate reducers

Older sulfate-rich groundwaters had the highest relative sequence abundance of sulfate reducers (Figure S10). *Desulfomicrobium* was the second most abundant genus occurring mainly in old waters, unlike *Desulfocapsa*, a clade known for sulfur disproportionation, which occurred on moderate to high relative abundance in many samples (66 of 108) across all water types. Sequences affiliating with the genus *Desulfomonas* known to perform sulfur reduction were also very widespread (found in 72 of 108 samples), except in young recharge waters. Overall, the aquifers apparently had a high functional redundancy harboring multiple sulfate-reducing populations. There were, however, some exceptions that were dominated by just one or few clades. In GW3104, GW378, GW969, GW218, GW298, GW117 and GW 155 we detected between two and four populations likely involved in sulfate reduction, while GW3002 harbored only one sulfate reducer affiliating with an unclassified genus of *Desulfobulbaceae*.

#### Sulfur oxidizers

Putative sulfur oxidizers were very widespread in the studied aquifers, including facultatively anaerobic *Sulfuricurvum sp*. which are known to oxidize sulfur with either oxygen or nitrate, and are also able to use hydrogen as an electron donor (*98*). Some *Thiobacillus sp*. can anaerobically oxidize pyrite using nitrite as electron acceptor (*99*), and may also use hydrogen, sulfur compounds or iron as electron donors (*100*). *Thiomicrorhabdus* have been shown to be versatile sulfur oxidizers in coal mine shafts (*101*). In two of the aquifers, they constituted 100 % of the sulfur oxidizers present (Fig S10, GW3026 and GW3004). *Sulfurimonas* is also known from subsurface ecosystems (*102*) and may oxidize sulfur compounds using nitrate, which was however found in few groundwater samples. *Sulfurimonadaceae* were mainly found in younger or older waters, while *Sulfuricellaceae* occurred in all groundwater types at low abundance. Sulfate-rich waters had a sulfur-isotopic signature indicating oxidation-derived sulfate (Fig 3B, D) of abiotic or biotic origin.

#### Other widespread clades

Aerobic *Nitrosopumilaceae* occurred mainly in young recharge waters, while an unclassified family within the *Woesearchaeales* was very abundant in all aquifers except those with older, mature waters. Metabolically versatile *Comamonadaceae* were found in all water ages at similar abundances, while *Hydrogenophilaceae* and *Pseudomonadaceae* preferentially occurred in sulfate-rich environments.

### Dismutases and catalases

Bacterial genera that have been shown to perform nitric oxide or chlorite dismutation were very widespread and often also very abundant in the studied aquifers. The genes coding for dismutases and catalases are widespread among microbial lineages (*56*, *103*–*105*). Groundwater aquifers can contain a high diversity of nitric oxide dismutation genes (*56*). Anoxic marine sediments were suggested to sustain aerobic microorganisms due to oxygen production via chlorite dismutation (*106*), and eukaryotic foraminifers that have a putative nitric oxide dismutase encoded in their genomes were shown to live in anoxic sediments (*107*).

### Contamination control

To avoid and monitor contamination we have included controls at each step of the experiments. We have sterilized the sampling gear after each well, we have included blind controls, i.e., samples replicating wells using encrypted sample names, and blank controls for cell counting and sequencing. We also checked the influence of sample handling and storage duration on the microbial community. We have not found any indications for contamination across the entire experiment. We have no indication that sample storage in Nalgene bottles had a potential enrichment effect, e.g., promoting growth of aerobic organisms after sampling. Both alpha diversity and beta diversity do not show any differences when grouped into categories reflecting storage time (Fig S11).

**Fig. S1.**
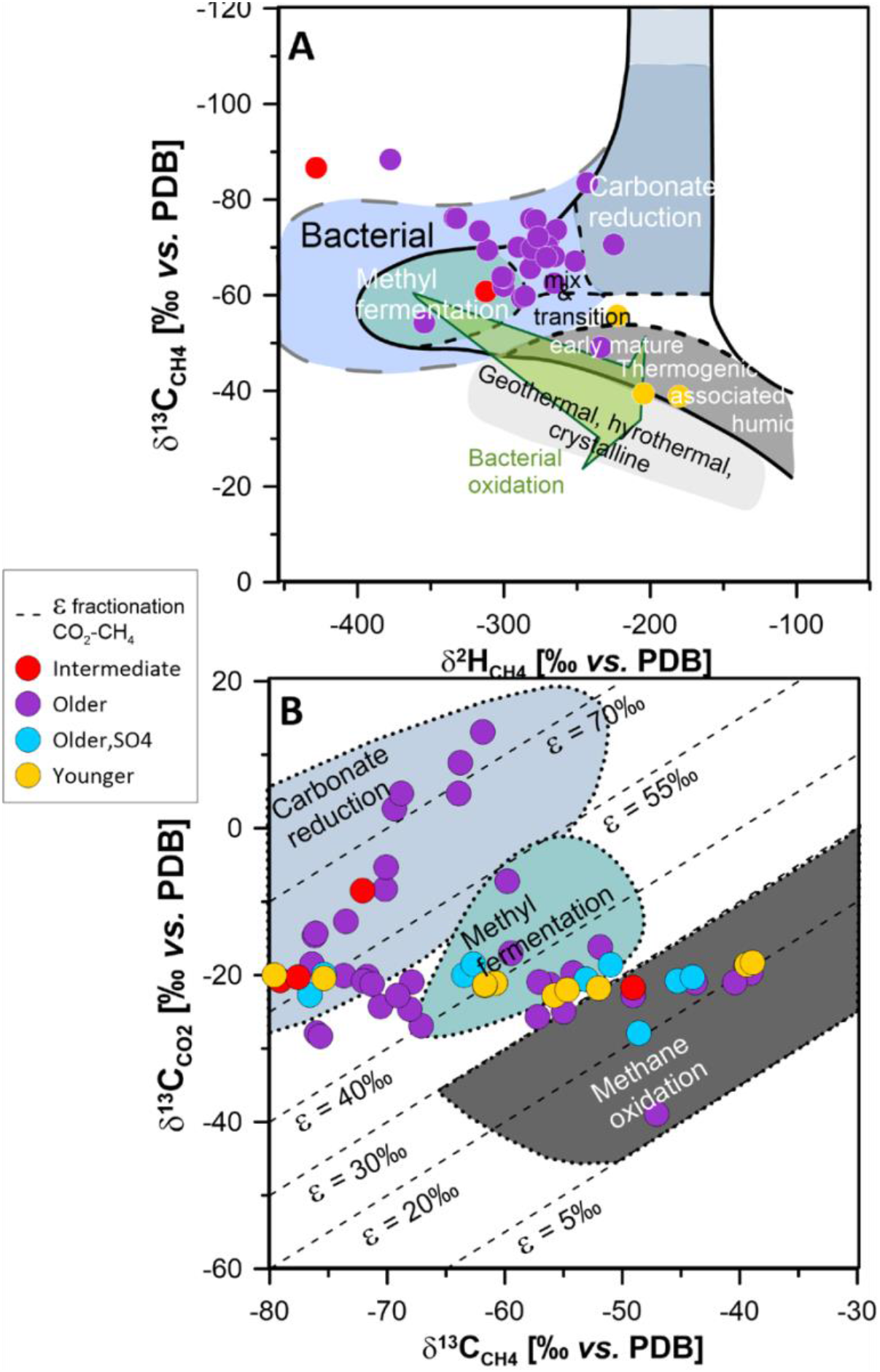
Methane carbon isotope geochemistry. Classification of microbial and thermogenic gas by combination of δ^13^C_CH4_ and δ^2^H_CH4_ (A) and δ^13^C_CO2_ and δ^13^C_CH4_ (B).

**Fig. S2.**
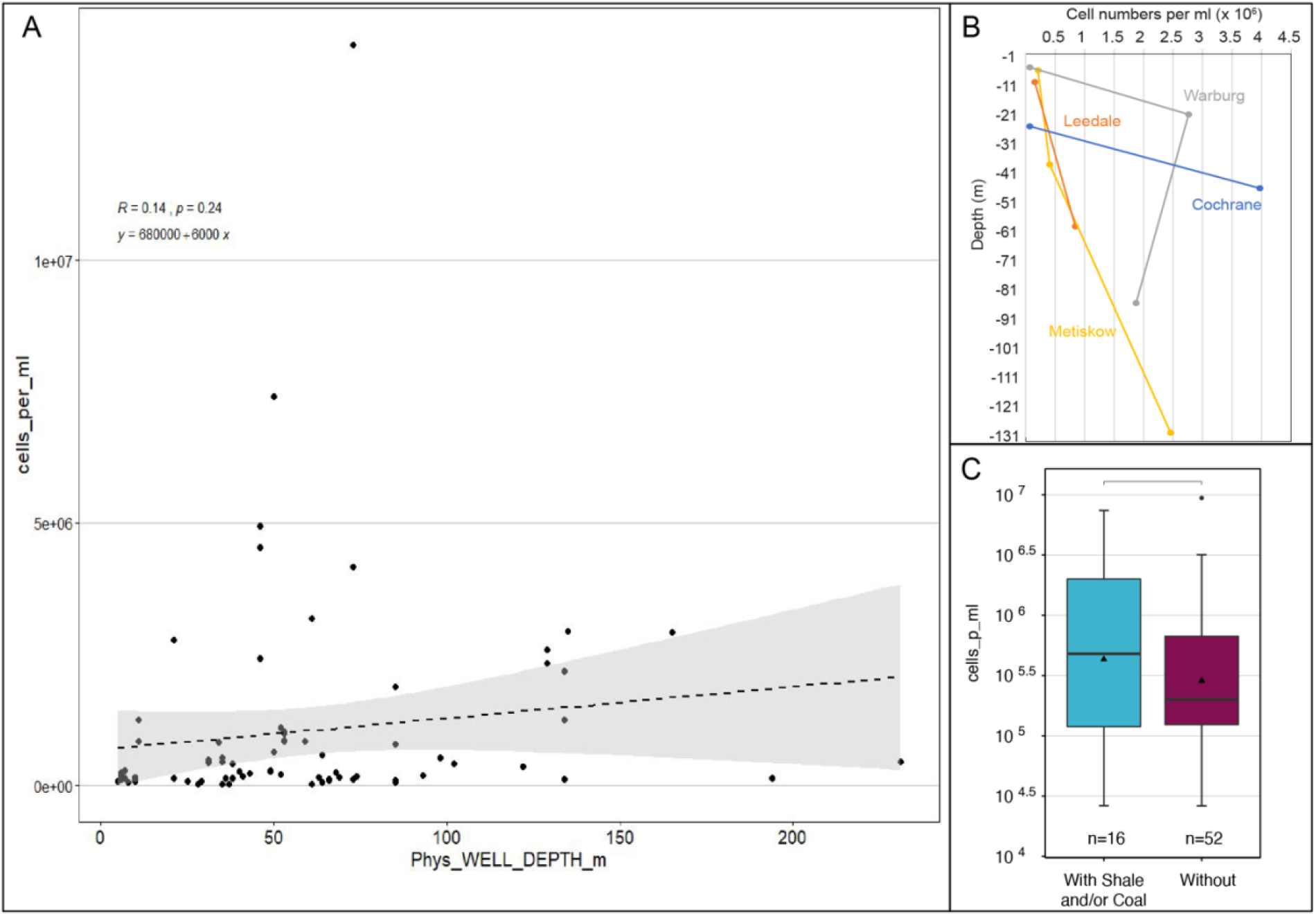
Microbial cell numbers. Cell abundance versus well depth across the entire dataset (A), and in selected aquifers for which multiple depth horizons were available to sample (B). Cell abundance in wells that were completed into formations with or without coal or shale or both. None of the trends are significant, however they indicate that cell abundance generally did not decrease with depth and may have been higher in energy-rich strata.

**Fig. S3.**
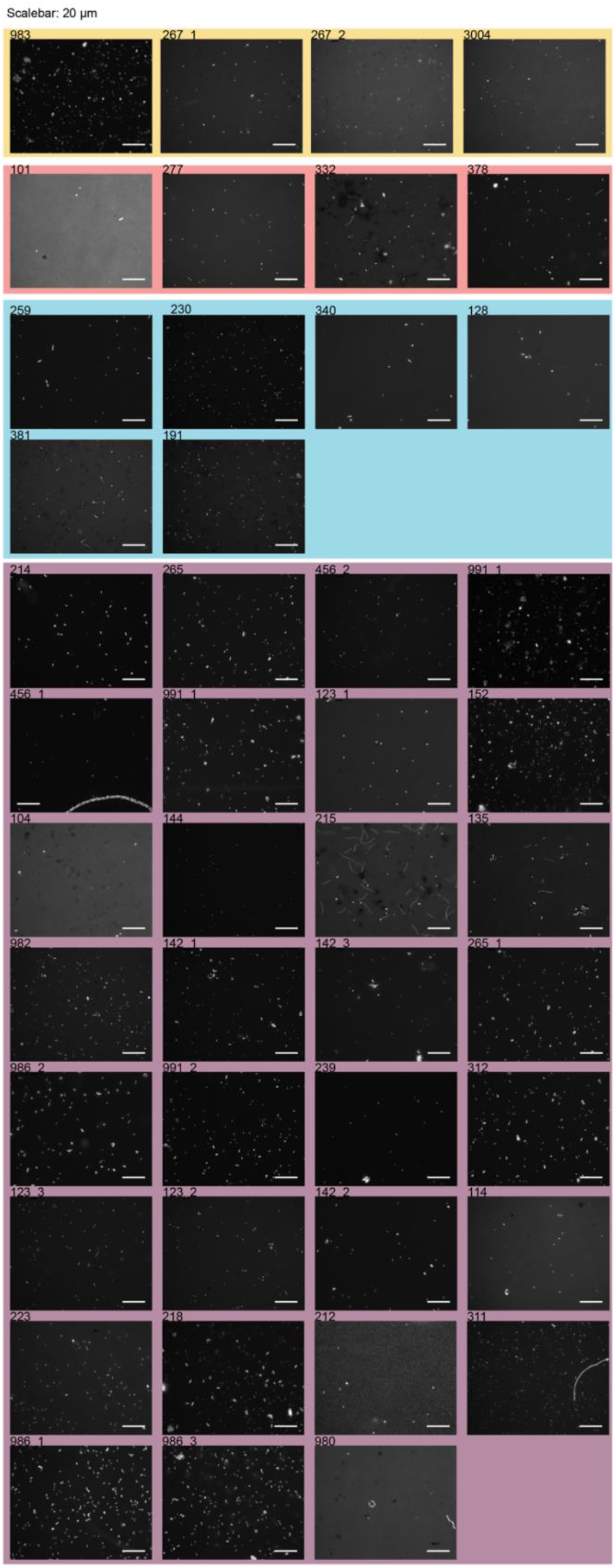
Fluorescence micrographs. Cell numbers and morphologies present in groundwater samples, as detected with the nucleic acid stain DAPI. The background color refers to the four water ages (yellow: younger, red: intermediate, blue: older with SO4, purple: older). Average cell size is similar in the different water types, which means that high cell number also mean high biomass. The scale bar is 20 μm in all pictures of the panel. A high-resolution .tif image is available in file Data S2.

**Fig. S4.**
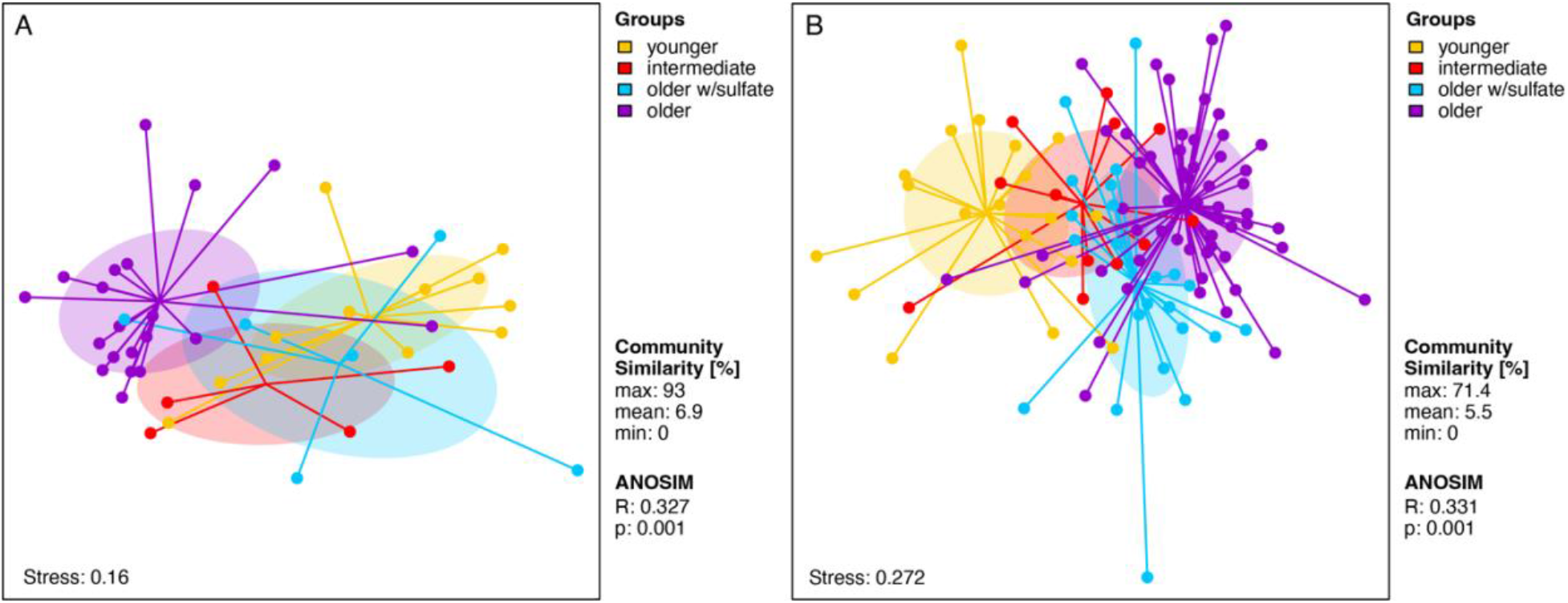
Microbial community shifts with water age. Archaeal (A) and bacterial (B) community structure shown as non-metric multidimensional ordinations based on community distance. Each circle is a sample. The further two samples are apart the more different are their underlying communities. Samples are grouped based on water types. Each sample is connected to the average weighted mean of within group distances (group centroid). Ellipses depict one standard deviation from the centroid. Nevertheless, we found a core community of around 1 % of all archaeal (32 ASV) and 3 % of all bacterial ASV (388 ASV) that occurred in all water types, suggesting the persistence of certain microbial populations in aquifers of different depths and ages (Figure S3).

**Fig. S5.**
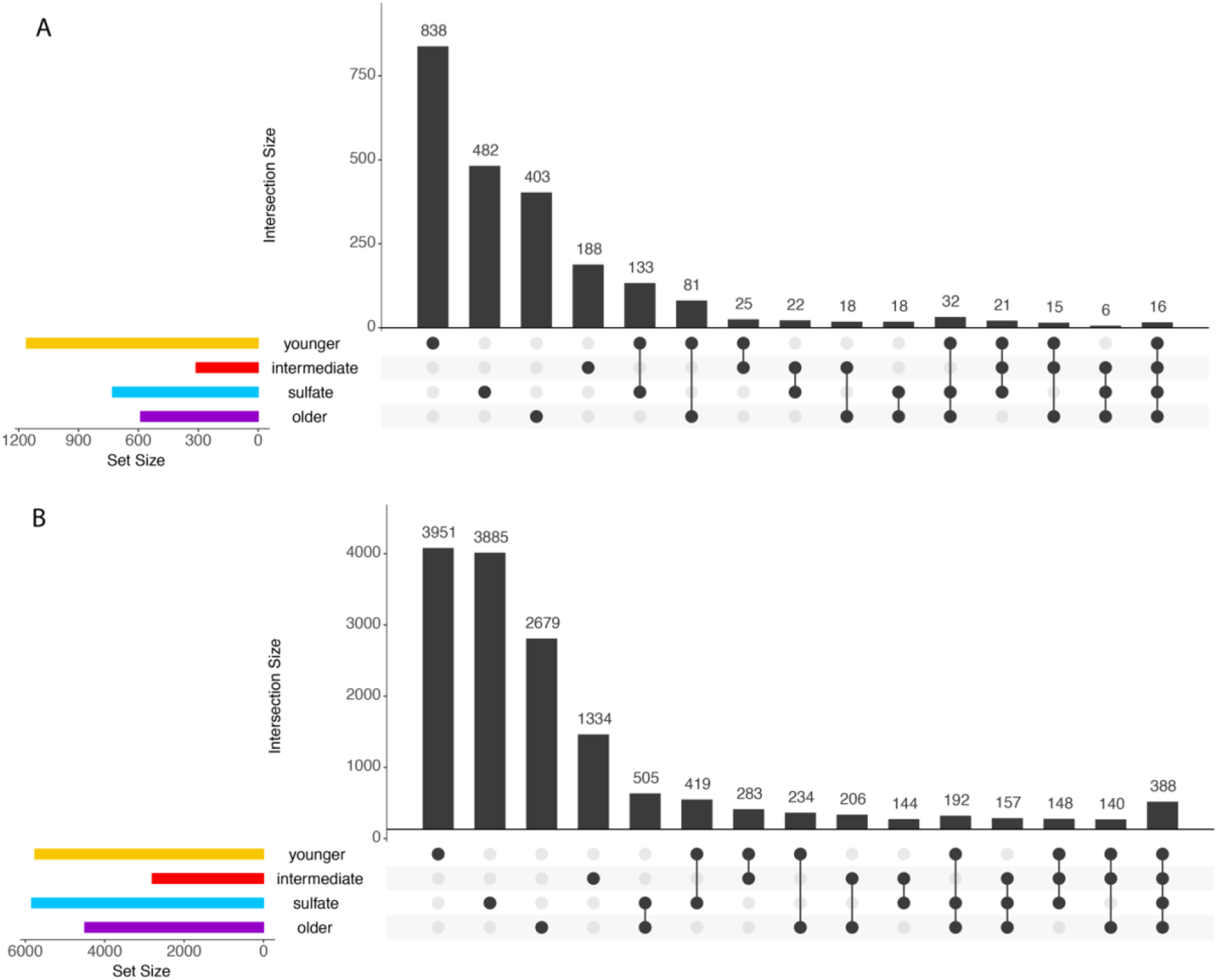
Number of unique and shared amplicon sequence variants. Upset plot showing archaeal (A) and bacterial (B) richness (number of amplicon sequence variants – ASV) that exclusively occur in the investigated water types and combination. The plot is to be read like a Venn diagram. The richness patterns are similar across domains with young (type I) waters showing the highest richness, followed by sulfate-rich intermediate (type III), old (type II) and sulfate-poor intermediate waters (type IV).

**Fig. S6.**
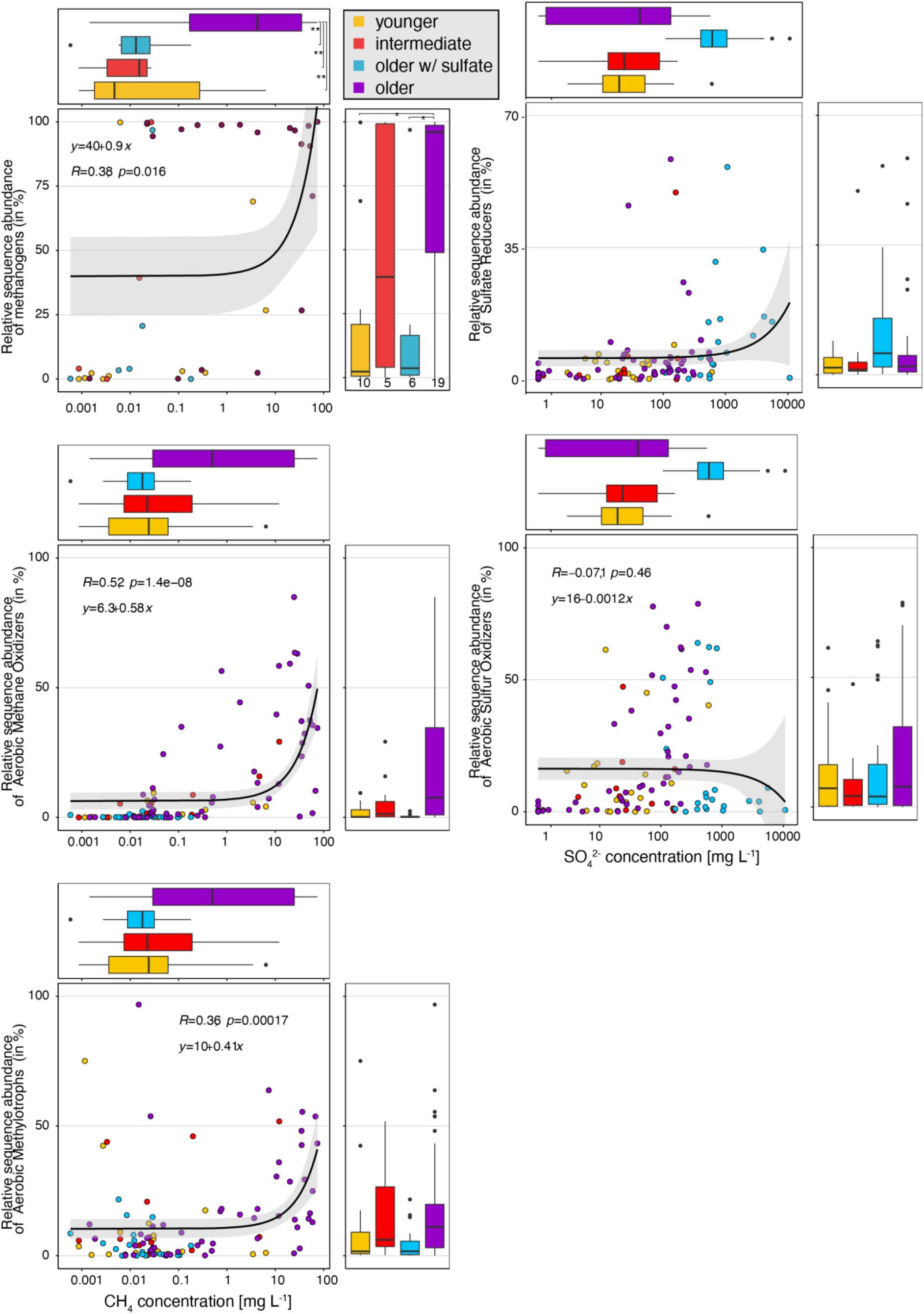
Methane and sulfur cycling microbes. Relative abundance of methanogens, aerobic methanotrophs and aerobic methylotrophs versus methane concentration (Left panel). There is a significant positive correlation with these methane-cycling clades with the concentration of methane. Sulfate reducers and sulfur oxidizers versus sulfate are shown in the right panel. Sulfate reducers tend to be abundant in sulfate-rich waters, while sulfide oxidizers are most abundant in waters with intermediate sulfate concentrations. Concentrations on the x-axis are shown on pseudo-log10 scale, so as to include the samples in which the compound was not detected (zero).

**Fig. S7.**
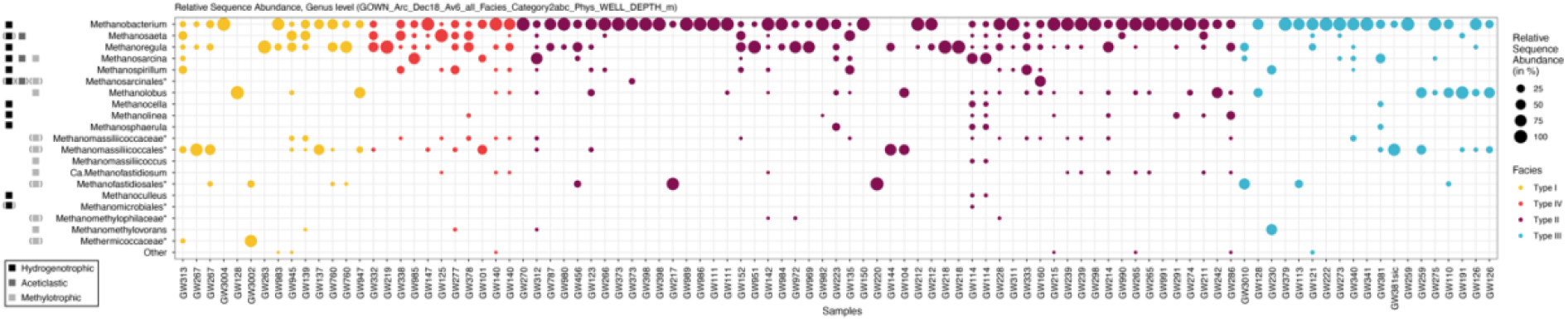
Methanogenic microbes. Relative sequence abundance of genus-level methanogenic clades based on 16S rRNA amplicon sequence variants (ASV). Clades marked by an asterisk represent genus level clades without cultured isolates, here the phylogenetic level with the closest cultured isolate is given. The different methanogenic capabilities hydrogenotrophic, acetoclastic and methylotrophic are shown as squares. Squares in brackets represent capabilities that are likely, yet not confirmed for the respective clade. Note: The shown relative abundance was calculated using only methanogenic clades and is thus not representative of these clades’ relative abundance in the whole community. Note to reviewers: This figure and the following figures will be available as high-resolution files at PANGAEA (submission in progress).

**Fig. S8.**
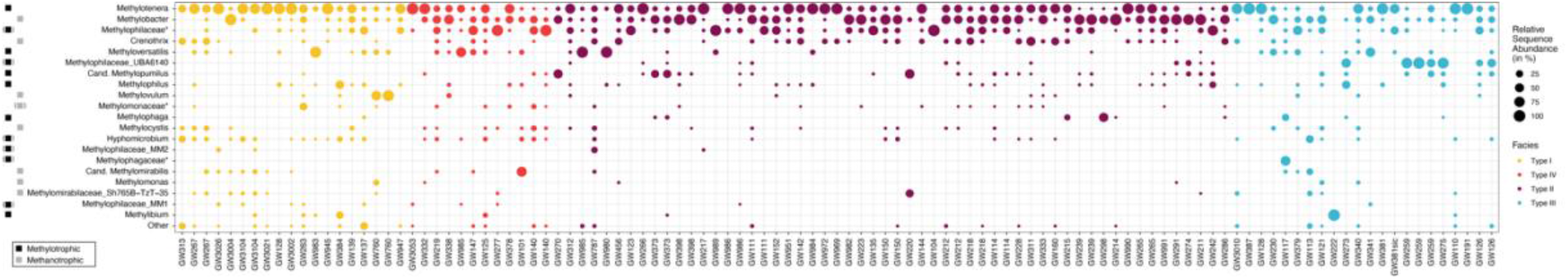
Methanotrophic microbes. Relative sequence abundance of genus-level aerobic methanotrophic clades based on 16S rRNA amplicon sequence variants (ASV). Clades marked by an asterisk represent genus level clades without cultured isolates, here the phylogenetic level with the closest cultured isolate is given. The different methanotrophic capabilities methanotrophic and methylotrophic are shown as squares. Squares in brackets represent capabilities that are likely, yet not confirmed for the respective clade. Note: The shown relative abundance was calculated using only methanotrophic clades and is thus not representative of these clades’ relative abundance in the whole community.

**Fig. S9.**
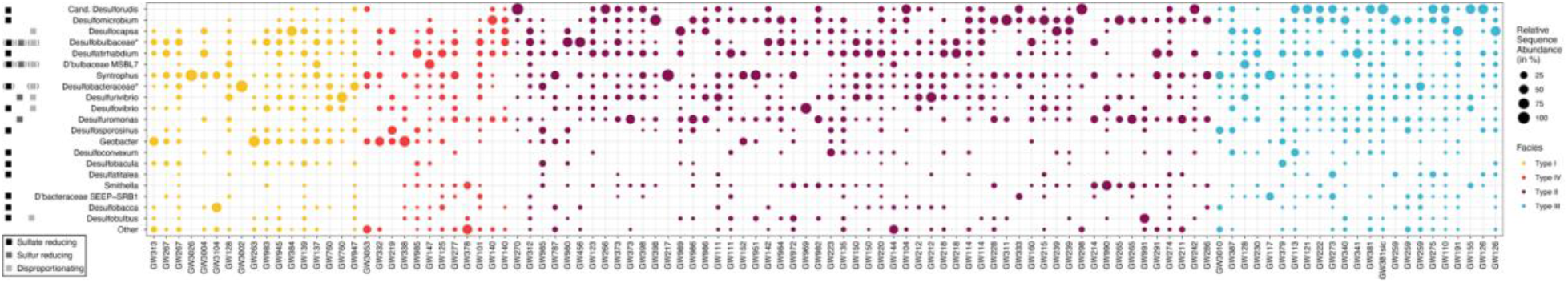
Sulfate-reducing microbes. Relative sequence abundance of genus-level sulfate-reducing clades based on 16S rRNA amplicon sequence variants (ASV). Clades marked by an asterisk represent genus level clades without cultured isolates, here the phylogenetic level with the closest cultured isolate is given. The different sulfur cycling capabilities are shown as squares. Squares in brackets represent capabilities that are likely, yet not confirmed for the respective clade. Note: The shown relative abundance was calculated using only sulfate-reducing clades and is thus not representative of these clades’ relative abundance in the whole community. Geobacter, Syntrophus and Smithella were included because they are close relatives and/or known to live syntrophically with hydrogen scavengers.

**Fig. S10.**
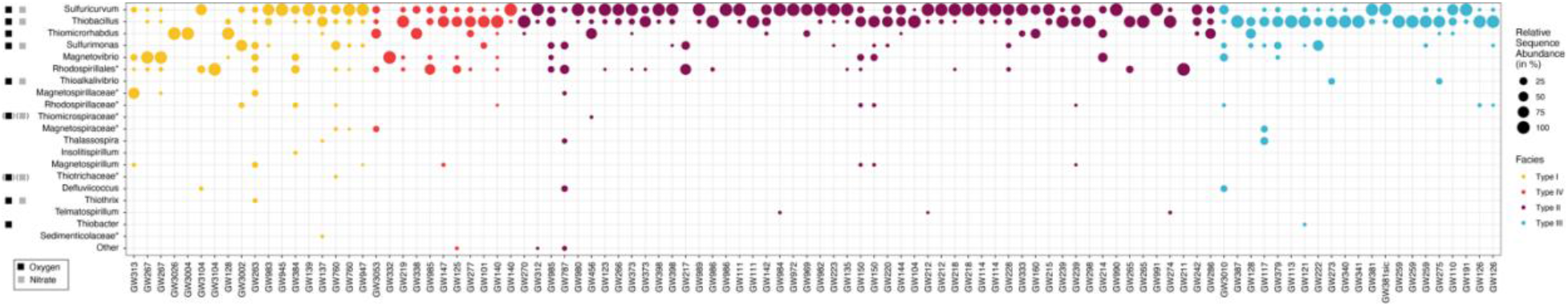
Sulfur-oxidizing microbes. Relative sequence abundance of genus-level sulfur-oxidizing clades based on 16S rRNA amplicon sequence variants (ASV). Clades marked by an asterisk represent genus level clades without cultured isolates, here the phylogenetic level with the closest cultured isolate is given. The different electron acceptor for sulfur oxidation are shown as squares. Squares in brackets represent capabilities that are likely, yet not confirmed for the respective clade. Note: The shown relative abundance was calculated using only thiotrophic clades and is thus not representative of these clades’ relative abundance in the whole community. Rhodospirillales and other non-sulfur-oxidizing clades were included because they are close relatives.

**Fig. S11.**
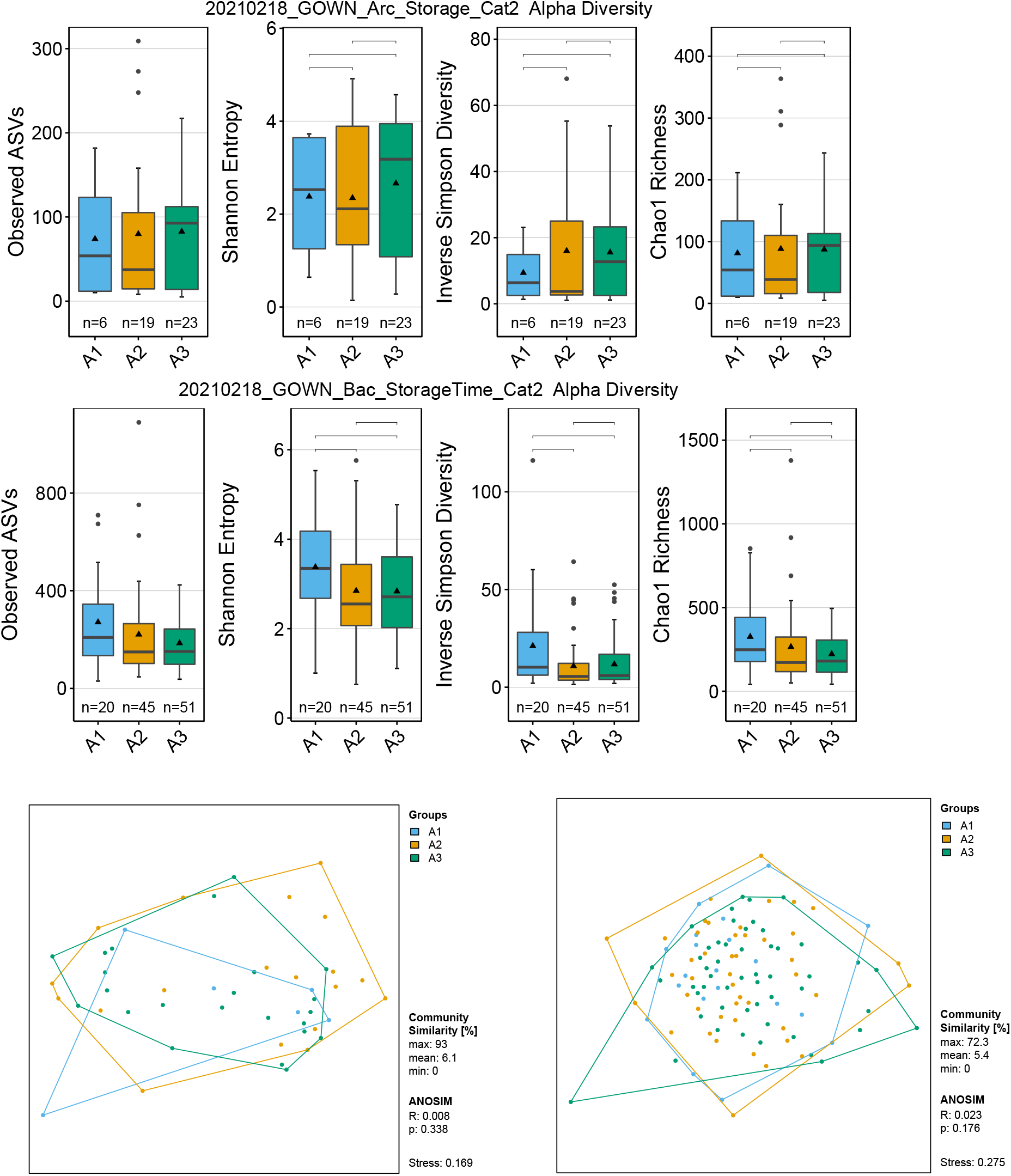
Community structure vs sample storage time. Archaeal and bacterial alpha and beta diversity grouped based on storage duration. Blue: 0-7 days, orange: 8-14 days, green: >14 days.

**Fig. S12.**
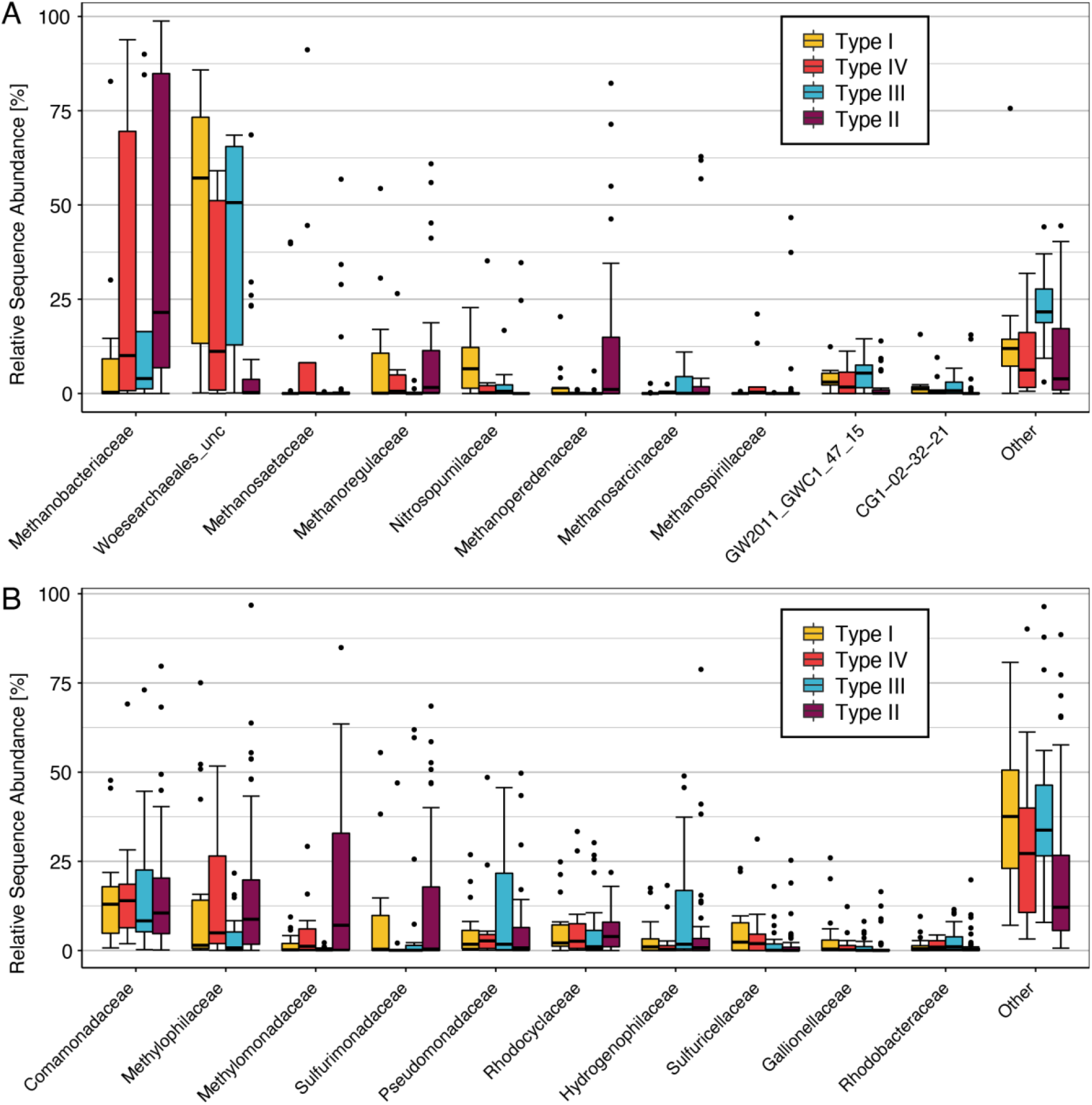
Most abundant family level microbial clades. Archaeal (A) and bacterial (B) relative sequence abundance of the 10 most abundant family-level clades based on 16S rRNA gene variable regions v6-v9 (archaea) or v3-v5 (bacteria). Boxplots represent the average relative sequence abundance of the given family-level clade for each water age. GW2011 and CG1 are clades within the *Woesearchaeales* and *Micrarchaeales* respectively.

**Table S1:** Depositional Environments and associated sediments in which the dedicated groundwater monitoring wells from the GOWN program are completed.

| Depositional Environments |  | # well | Formation |
| --- | --- | --- | --- |
| **Neogene-Quaternary surficial deposits** |  |  |  |
| Buried Channel* (a-preglacial fluvial incision, b-glacialfluvial incision, c-unknown) |  | 16 | Medicine Hat Valley<sup>a</sup>, Lethbridge Valley<sup>a</sup>, Keho Valley, Irvine Valley, Sibbald Valley, Vermilion Valley, Bronson Valley |
| Undifferentiated/Glacial tills |  | 22 |  |
| **Upper Cretaceous to Paleocene** |  |  |  |
| Coarse sandy clastic deposits (terrestrial env.) |  | 3<br>7<br>4<br>9 | Milk River,<br>Belly River Group (e.g. Oldman),<br>Horseshoe Canyon,<br>Scollard/Paskapoo, |
| Organic-rich/coal seams (terrestrial env.) |  | 1<br>6<br>2 | Belly River,<br>Horseshoe Canyon,<br>Paskapoo |
| Fined-grained deposits<br>(*shale, siltstone, mudstone*) | Marine env./transgr. | 4<br>2 | Bearpaw,<br>Loon River |
|  | Terrestrial env. | 1<br>3<br>14 | Belly River Group (e.g., Bulwark, Foremost),<br>Horseshoe Canyon,<br>Scollard/Paskapoo |
|  | Undifferentiated | 1 |  |
| *\*Buried valleys are referring to valleys that are incised into bedrock and are buried by till.* |  |  |  |

**Table S2:** Archaeal diversity Indices. Apart from total reads the diversity indices were calculated using iterated subsampling of 1008 reads.

| Sample | Total reads | Observed ASV | Estimated Richness (Chao1) | Inverse Simpson Diversity | Shannon Entropy |
| --- | --- | --- | --- | --- | --- |
| GW980_BE128_S99_Av6 | 1230 | 3 | 3 | 1 | 0.2 |
| GW239_BE017_S57_Av6 | 2380 | 5 | 6 | 1 | 0.5 |
| GW265_BE060_S123_Av6 | 2274 | 5 | 6 | 1 | 0.5 |
| GW984_BE083_S101_Av6 | 7891 | 5 | 5 | 3 | 1.1 |
| GW239_BE018_S66_Av6 | 2492 | 6 | 6 | 1 | 0.6 |
| GW991_BE075_S139_Av6 | 1447 | 6 | 6 | 2 | 1 |
| GW140_BE061_S75_Av6 | 13217 | 7 | 6 | 1 | 0.3 |
| GW340_BE107_S118_Av6 | 1274 | 7 | 7 | 1 | 0.6 |
| GW982_BE129_S135_Av6 | 14249 | 7 | 7 | 1 | 0.3 |
| GW983_BE084_S60_Av6 | 21876 | 7 | 13 | 1 | 0.5 |
| GW991_BE077_S141_Av6 | 2344 | 7 | 7 | 2 | 1.1 |
| GW239_BE019_S80_Av6 | 8773 | 8 | 10 | 1 | 0.6 |
| GW991_BE076_S140_Av6 | 5401 | 8 | 9 | 2 | 1.1 |
| GW111_BE008_S100_Av6 | 1196 | 9 | 9 | 3 | 1.3 |
| GW311_BE116_S133_Av6 | 1147 | 9 | 9 | 4 | 1.5 |
| GW275_BE137_S88_Av6 | 2356 | 10 | 10 | 1 | 0.6 |
| GW140_BE131_S70_Av6 | 9437 | 11 | 12 | 2 | 0.8 |
| GW211_BE054_S116_Av6 | 1442 | 11 | 11 | 3 | 1.4 |
| GW239_BE082_S121_Av6 | 14576 | 11 | 13 | 2 | 0.7 |
| GW291_BE143_S144_Av6 | 3692 | 11 | 11 | 3 | 1.5 |
| GW333_BE025_S81_Av6 | 1018 | 11 | 11 | 4 | 1.7 |
| GW142_BE097_S108_Av6 | 2077 | 12 | 12 | 2 | 1.1 |
| GW228_BE070_S31_Av6 | 10464 | 12 | 14 | 2 | 1.2 |
| GW398_BE127_S93_Av6 | 1946 | 12 | 13 | 3 | 1.4 |
| GW142_BE099_S97_Av6 | 9481 | 13 | 18 | 2 | 1.1 |
| GW152_BE089_S143_Av6 | 6126 | 14 | 19 | 3 | 1.5 |
| GW338_BE041_S95_Av6 | 1342 | 14 | 14 | 5 | 1.7 |
| GW135_BE125_S128_Av6 | 33349 | 16 | 18 | 2 | 1.2 |
| GW160_BE026_S115_Av6 | 7174 | 16 | 17 | 4 | 1.6 |
| GW125_BE106_S113_Av6 | 1789 | 17 | 17 | 1 | 0.7 |
| GW160_BE026_S160_Av6 | 4228 | 18 | 22 | 3 | 1.6 |
| GW214_BE005_S79_Av6 | 1599 | 18 | 18 | 5 | 2 |
| GW951_BE016_S7_Av6 | 5182 | 19 | 23 | 4 | 1.6 |
| GW972_BE057_S17_Av6 | 5834 | 19 | 21 | 3 | 1.6 |
| GW114_BE118_S106_Av6 | 1151 | 20 | 20 | 5 | 2.1 |
| GW456_BE068_S102_Av6 | 1976 | 26 | 27 | 3 | 1.8 |
| GW114_BE029_S85_Av6 | 10665 | 27 | 30 | 5 | 2.1 |
| GW139_BE095_S112_Av6 | 1173 | 27 | 27 | 6 | 2.2 |
| GW218_BE031_S22_Av6 | 1728 | 28 | 28 | 5 | 2.2 |
| GW312_BE074_S76_Av6 | 11542 | 30 | 41 | 3 | 1.7 |
| GW456_BE067_S91_Av6 | 4723 | 36 | 37 | 3 | 1.8 |
| GW456_BE069_S83_Av6 | 4937 | 36 | 38 | 3 | 1.8 |
| GW147_BE100_S42_Av6 | 1156 | 38 | 38 | 3 | 1.9 |
| GW110_BE003_S87_Av6 | 1482 | 44 | 44 | 19 | 3.3 |
| GW137_BE124_S96_Av6 | 2191 | 49 | 50 | 5 | 2.4 |
| GW760_BE104_S18_Av6 | 2426 | 56 | 61 | 3 | 2.1 |
| GW3004_BE065_S29_Av6 | 1008 | 69 | 72 | 29 | 3.7 |
| GW220_BE048_S27_Av6 | 1038 | 71 | 71 | 23 | 3.7 |
| GW985_BE111_S62_Av6 | 2446 | 79 | 82 | 13 | 3.2 |
| GW144_BE085_S90_Av6 | 2305 | 87 | 92 | 17 | 3.6 |
| GW117_BE052_S34_Av6 | 2187 | 90 | 93 | 23 | 3.7 |
| GW121_BE034_S69_Av6 | 2079 | 92 | 96 | 18 | 3.6 |
| GW379_BE108_S122_Av6 | 2032 | 94 | 98 | 25 | 3.9 |
| GW381_BE138_S73_Av6 | 2983 | 100 | 108 | 29 | 3.9 |
| GW3104_BE079_S46_Av6 | 10755 | 102 | 107 | 40 | 4.1 |
| GW277_BE093_S65_Av6 | 2219 | 103 | 110 | 21 | 3.7 |
| GW3053_BE117_S36_Av6 | 2493 | 112 | 115 | 21 | 3.9 |
| GW3050_BE119_S10_Av6 | 4432 | 116 | 143 | 8 | 3 |
| GW760_BE037_S30_Av6 | 5868 | 116 | 139 | 17 | 3.5 |
| GW101_BE140_S64_Av6 | 1559 | 136 | 143 | 45 | 4.4 |
| GW3002_BE130_S32_Av6 | 3977 | 144 | 159 | 49 | 4.4 |
| GW313_BE115_S40_Av6 | 9625 | 146 | 176 | 26 | 4 |
| GW787_BE062_S15_Av6 | 25568 | 150 | 201 | 9 | 3.6 |
| GW3026_BE014_S45_Av6 | 13158 | 171 | 201 | 51 | 4.5 |
| GW267_BE059_S43_Av6 | 8137 | 182 | 224 | 46 | 4.5 |
| GW267_BE134_S68_Av6 | 9160 | 222 | 296 | 26 | 4.4 |
| GW3010_BE006_S5_Av6 | 14862 | 242 | 336 | 65 | 4.8 |

**Table S3:** Bacterial Diversity Indices. Apart from total reads the diversity indices were calculated using iterated subsampling of 6796 reads.

| Sample | Reads | Observed<br>ASV | Estimated<br>Richness<br>(Chao1) | Inverse<br>Simpson<br>Diversity | Shannon<br>Entropy |
| --- | --- | --- | --- | --- | --- |
| GW969_BE007_S119 | 29092 | 30 | 35 | 2 | 1 |
| GW986_BE121_S147 | 30001 | 34 | 41 | 5 | 2 |
| GW333_BE025_S158 | 29663 | 35 | 42 | 4 | 1.6 |
| GW986_BE123_S148 | 41177 | 46 | 56 | 5 | 2 |
| GW982_BE129_S135 | 22379 | 47 | 48 | 5 | 2.1 |
| GW218_BE120_S145 | 55447 | 48 | 59 | 3 | 1.5 |
| GW114_BE118_S106 | 14407 | 50 | 55 | 4 | 2 |
| GW333_BE025_S81 | 48144 | 50 | 67 | 4 | 1.6 |
| GW986_BE122_S142 | 39708 | 50 | 59 | 6 | 2.1 |
| GW212_BE004_S103 | 16577 | 55 | 59 | 3 | 1.8 |
| GW298_BE039_S107 | 6796 | 55 | 55 | 5 | 2.2 |
| GW123_BE081_S105 | 5847 | 58 | 58 | 3 | 1.7 |
| GW142_BE097_S108 | 12367 | 60 | 66 | 6 | 2.2 |
| GW152_BE089_S143 | 49272 | 63 | 81 | 4 | 1.7 |
| GW214_BE005_S79 | 47318 | 63 | 89 | 1 | 0.8 |
| GW265_BE132_S134 | 41067 | 64 | 75 | 3 | 1.8 |
| GW456_BE068_S102 | 20222 | 65 | 72 | 3 | 1.6 |
| GW341_BE043_S44 | 9334 | 70 | 74 | 7 | 2.5 |
| GW991_BE075_S139 | 54198 | 70 | 87 | 3 | 1.7 |
| GW991_BE077_S141 | 43354 | 72 | 86 | 3 | 1.7 |
| GW991_BE076_S140 | 47247 | 73 | 94 | 3 | 1.8 |
| GW378_BE112_S137 | 38144 | 74 | 86 | 4 | 2.2 |
| GW951_BE016_S7 | 58291 | 74 | 107 | 4 | 1.7 |
| GW266_BE133_S127 | 36226 | 77 | 89 | 4 | 2.1 |
| GW259_BE002_S38 | 16687 | 78 | 87 | 4 | 2 |
| GW951_BE016_S155 | 61652 | 81 | 113 | 3 | 1.6 |
| GW456_BE067_S91 | 39001 | 82 | 96 | 3 | 1.6 |
| GW265_BE060_S123 | 31431 | 85 | 95 | 4 | 2.1 |
| GW990_BE053_S125 | 29414 | 87 | 92 | 4 | 2.2 |
| GW111_BE008_S100 | 41375 | 88 | 108 | 3 | 1.9 |
| GW140_BE131_S70 | 28147 | 88 | 109 | 4 | 1.8 |
| GW214_BE005_S78 | 33234 | 89 | 119 | 3 | 1.9 |
| GW212_BE113_S126 | 29458 | 95 | 108 | 12 | 2.9 |
| GW291_BE143_S144 | 39037 | 95 | 114 | 5 | 2.2 |
| GW223_BE110_S146 | 52949 | 96 | 117 | 7 | 2.6 |
| GW3053_BE117_S36 | 41553 | 96 | 145 | 3 | 1.5 |
| GW128_BE103_S120 | 43967 | 97 | 116 | 7 | 2.5 |
| GW398_BE009_S13 | 52126 | 100 | 111 | 7 | 2.6 |
| GW270_BE030_S89 | 34414 | 102 | 117 | 3 | 2 |
| GW142_BE099_S97 | 28260 | 106 | 121 | 6 | 2.5 |
| GW211_BE054_S116 | 55296 | 106 | 135 | 3 | 2 |
| GW456_BE069_S83 | 59936 | 107 | 135 | 3 | 1.9 |
| GW104_BE087_S56 | 18687 | 108 | 124 | 2 | 1.4 |
| GW311_BE116_S133 | 41757 | 108 | 137 | 6 | 2.3 |
| GW986_BE011_S23 | 49289 | 109 | 119 | 3 | 2.2 |
| GW239_BE018_S66 | 41876 | 114 | 137 | 3 | 1.9 |
| GW239_BE019_S80 | 49365 | 114 | 143 | 3 | 1.9 |
| GW760_BE104_S18 | 43064 | 116 | 143 | 3 | 2.1 |
| GW128_BE015_S156 | 56170 | 119 | 149 | 5 | 2.2 |
| GW398_BE049_S58 | 59118 | 119 | 138 | 8 | 2.7 |
| GW128_BE015_S24 | 51337 | 122 | 154 | 3 | 2 |
| GW218_BE031_S22 | 48430 | 124 | 161 | 9 | 2.8 |
| GW986_BE013_S54 | 48306 | 126 | 147 | 6 | 2.7 |
| GW398_BE009_S157 | 76734 | 129 | 179 | 2 | 1.6 |
| GW160_BE026_S115 | 44438 | 130 | 164 | 5 | 2.3 |
| GW989_BE055_S39 | 22325 | 130 | 142 | 8 | 3 |
| GW972_BE057_S17 | 50475 | 131 | 168 | 4 | 2.2 |
| GW3002_BE130_S32 | 48306 | 134 | 178 | 2 | 1.6 |
| GW986_BE012_S4 | 54861 | 134 | 153 | 3 | 2.4 |
| GW980_BE128_S99 | 21835 | 141 | 153 | 10 | 3.3 |
| GW111_BE141_S129 | 28938 | 144 | 170 | 4 | 2.5 |
| GW984_BE083_S101 | 18674 | 146 | 154 | 21 | 3.6 |
| GW312_BE074_S76 | 30401 | 147 | 177 | 6 | 2.7 |
| GW229_BE071_S136 | 36960 | 148 | 169 | 7 | 2.9 |
| GW373_BE042_S19 | 53058 | 155 | 195 | 7 | 2.8 |
| GW126_BE001_S63 | 35965 | 156 | 187 | 8 | 2.9 |
| GW338_BE041_S95 | 36545 | 156 | 173 | 13 | 3.5 |
| GW398_BE127_S93 | 45225 | 157 | 189 | 7 | 2.9 |
| GW160_BE026_S160 | 41919 | 159 | 195 | 5 | 2.5 |
| GW110_BE003_S154 | 38492 | 160 | 198 | 7 | 2.7 |
| GW114_BE029_S85 | 58357 | 161 | 202 | 5 | 2.6 |
| GW398_BE009_S14 | 72900 | 163 | 198 | 3 | 2.2 |
| GW123_BE091_S110 | 40374 | 170 | 209 | 4 | 2.4 |
| GW239_BE017_S57 | 55165 | 170 | 215 | 6 | 2.7 |
| GW222_BE035_S59 | 36223 | 172 | 196 | 15 | 3.3 |
| GW275_BE137_S88 | 26762 | 173 | 188 | 10 | 3.4 |
| GW332_BE101_S131 | 26738 | 177 | 214 | 5 | 2.7 |
| GW220_BE048_S27 | 50157 | 178 | 241 | 5 | 2.4 |
| GW230_BE072_S132 | 41939 | 180 | 209 | 6 | 2.8 |
| GW273_BE136_S84 | 71384 | 184 | 224 | 8 | 3.3 |
| GW274_BE135_S104 | 14520 | 191 | 209 | 29 | 4 |
| GW373_BE105_S114 | 44794 | 200 | 244 | 20 | 3.9 |
| GW239_BE082_S121 | 67112 | 202 | 256 | 7 | 3.1 |
| GW277_BE093_S65 | 50038 | 202 | 265 | 6 | 2.7 |
| GW985_BE111_S62 | 33046 | 213 | 245 | 18 | 3.7 |
| GW228_BE070_S31 | 62585 | 215 | 264 | 11 | 3.3 |
| GW985_BE010_S1 | 49934 | 218 | 246 | 22 | 3.9 |
| GW947_BE078_S77 | 32434 | 226 | 266 | 7 | 3.3 |
| GW144_BE085_S90 | 26465 | 227 | 291 | 18 | 3.6 |
| GW340_BE107_S118 | 46274 | 229 | 264 | 5 | 3 |
| GW3050_BE119_S10 | 47470 | 231 | 307 | 2 | 1.9 |
| GW150_BE023_S67 | 47168 | 243 | 274 | 34 | 4.2 |
| GW150_BE024_S92 | 43201 | 244 | 281 | 25 | 4 |
| GW259_BE126_S25 | 59605 | 245 | 308 | 9 | 3.3 |
| GW387_BE051_S130 | 47906 | 254 | 344 | 5 | 3 |
| GW126_BE142_S28 | 28152 | 255 | 306 | 15 | 3.6 |
| GW983_BE084_S60 | 105629 | 269 | 335 | 19 | 3.8 |
| GW121_BE034_S69 | 48173 | 276 | 361 | 5 | 3.1 |
| GW150_BE066_S55 | 55119 | 280 | 342 | 31 | 4.1 |
| GW140_BE061_S75 | 34876 | 283 | 316 | 47 | 4.5 |
| GW135_BE125_S128 | 50999 | 284 | 353 | 9 | 3.4 |
| GW3004_BE065_S29 | 58075 | 288 | 392 | 4 | 2.6 |
| GW381_BE138_S73 | 56973 | 295 | 380 | 19 | 3.7 |
| GW760_BE037_S30 | 62880 | 306 | 403 | 13 | 3.6 |
| GW121_BE033_S8 | 40155 | 324 | 389 | 18 | 4.1 |
| GW267_BE059_S43 | 22892 | 325 | 389 | 16 | 4 |
| GW101_BE140_S64 | 36913 | 361 | 457 | 12 | 3.9 |
| GW147_BE100_S42 | 32073 | 362 | 449 | 12 | 3.8 |
| GW139_BE095_S112 | 45209 | 378 | 463 | 22 | 4.2 |
| GW379_BE108_S122 | 38837 | 382 | 452 | 52 | 4.8 |
| GW219_BE032_S35 | 48775 | 383 | 471 | 49 | 4.6 |
| GW3026_BE014_S45 | 48159 | 385 | 482 | 6 | 3.4 |
| GW125_BE106_S113 | 46436 | 390 | 469 | 45 | 4.7 |
| GW313_BE115_S40 | 21882 | 390 | 442 | 44 | 4.7 |
| GW137_BE124_S96 | 36213 | 401 | 486 | 43 | 4.7 |
| GW3104_BE079_S46 | 41735 | 409 | 473 | 19 | 4.2 |
| GW384_BE144_S61 | 51958 | 418 | 523 | 44 | 4.6 |
| GW110_BE003_S87 | 48814 | 422 | 520 | 14 | 3.9 |
| GW263_BE056_S50 | 16383 | 430 | 483 | 61 | 4.9 |
| GW117_BE052_S34 | 49680 | 492 | 623 | 15 | 4.3 |
| GW787_BE062_S15 | 43232 | 686 | 840 | 116 | 5.5 |
| GW267_BE134_S68 | 40318 | 718 | 911 | 29 | 5.1 |
| GW3010_BE006_S5 | 75554 | 1038 | 1359 | 63 | 5.7 |

**Table S4:** Environmental fitting using archaeal ASV data.

| Parameter | NMDS1 | NMDS2 | Envfit<br>Vectors | Envfit<br>Vector<br>s p-<br>value | Significanc<br>e Level |
| --- | --- | --- | --- | --- | --- |
|  | 0.89192491 |  | 0.21676002 |  |  |
| Gas_METHANE_ugL | 1 | 0.45218354 | 7 | 0.001 | *** |
|  | 0.36480317 | 0.93108465 | 0.21729311 |  |  |
| Chem_HCO3_calc_mgL | 9 | 8 | 3 | 0.001 | *** |
|  | 0.94863432 | 0.31637464 | 0.24375658 |  |  |
| Gas_CO2_mgL | 6 | 5 | 1 | 0.001 | *** |
|  | - | 0.47249867 | 0.24505098 |  |  |
| Chem_Mg_dis_mgL | 0.88133138 | 5 | 7 | 0.001 | *** |
|  | 0.44663784 | 0.89471483 | 0.25902174 |  |  |
| Chem_Alkalinity_CACO3_mgL | 8 | 3 | 4 | 0.001 | *** |
|  | 0.95556932 |  | 0.33966495 |  |  |
| Chem_Hardness_CACO3_mgL | 5 | 0.29476646 | 2 | 0.001 | *** |
|  | 0.98452625 | 0.17523714 | 0.35946057 |  |  |
| Chem_PH_LAB | 4 | 2 | 6 | 0.001 | *** |
|  |  | - | 0.42938118 |  |  |
| Phys_WELL_DEPTH_m | 0.99355088 | 0.11338716 | 8 | 0.001 | *** |
|  | 0.99922496 | 0.03936334 | 0.48589314 |  |  |
| Chem_Ca_dis_mgL | 3 | 1 | 2 | 0.001 | *** |
|  | 0.50697683 | 0.86195967 | 0.15523106 |  |  |
| Gas_dDH2O | 6 | 9 | 1 | 0.004 | *** |
|  | 0.66456956 | 0.74722639 |  |  |  |
| Chem_F_dis_mgL | 7 | 8 | 0.18659805 | 0.004 | *** |
|  | 0.72306762 | 0.69077724 | 0.17687385 |  |  |
| Chem_N2_calc_mgL | 7 | 8 | 3 | 0.005 | *** |
|  | 0.61255225 | 0.79043009 |  |  |  |
| Gas_d13CDIC | 2 | 7 | 0.14362687 | 0.008 | *** |
|  | 0.55714630 | 0.83041435 | 0.14694817 |  |  |
| Gas_d18OH2O | 3 | 2 | 5 | 0.008 | *** |
|  | 0.48154032 | - | 0.13312107 |  |  |
| Chem_Silica_soluble_reactive_mgL | 1 | 0.87642394 | 1 | 0.011 | ** |
|  | 0.70376252 | 0.71043529 | 0.13656424 |  |  |
| Chem_SO4_dis_mgL | 9 | 1 | 5 | 0.011 | ** |
|  | 0.95322931 | - | 0.14142485 |  |  |
| Chem_K_dis_mgL | 1 | 0.30224804 | 5 | 0.011 | ** |
|  | 0.16171075 |  | 0.13002778 |  |  |
| Phys_Lat | 7 | -0.9868382 | 2 | 0.017 | ** |
|  | 0.43165145 | 0.90204047 | 0.11339267 |  |  |
| Chem_tot_dis_solids_calc_mgL | 2 | 8 | 8 | 0.018 | ** |
|  | 0.61062601 | 0.79191910 |  |  |  |
| Gas_ETHANE_ugL | 7 | 4 | 0.11720432 | 0.022 | ** |
|  | 0.99964984 | 0.02646094 | 0.11043314 |  |  |
| Chem_P_mgL | 8 | 1 | 3 | 0.026 | ** |
|  | 0.07998107 | 0.99679638 |  |  |  |
| Chem_Na_dis_mgL | 4 | 2 | 0.09802801 | 0.035 | ** |
| Chem_spec_conductance_LAB_u | 0.23125940 | 0.97289212 | 0.09389320 |  |  |
| Scm | 5 | 5 | 6 | 0.039 | ** |
|  | 0.66282701 | 0.74877256 | 0.08812248 |  |  |
| Chem_Cl_dis_mgL | 5 | 1 | 4 | 0.04 | ** |
|  | 0.28633119 | 0.95813070 | 0.09500692 |  |  |
| Phys_Elevation_mAMSL | 2 | 5 | 3 | 0.044 | ** |
|  | 0.71678325 | - | 0.04189348 |  |  |
| Gas_N2_mgL | 6 | 0.69729604 | 8 | 0.247 |  |
|  | 0.88178835 | - | 0.03112327 |  |  |
| Gas_O2_mgL | 6 | 0.47164531 | 7 | 0.359 |  |
|  | - |  | 0.02687891 |  |  |
| Sampling_Date | 0.52445527 | -0.851438 | 9 | 0.417 |  |
|  | 0.70892727 | 0.70528158 | 0.01280455 |  |  |
| Phys_Lon | 5 | 9 | 3 | 0.65 |  |
|  |  | 0.28164867 | 0.00020340 |  |  |
| Gas_PROPANE_ugL | 0.9595176 | 4 | 9 | 0.993 |  |

**Table S5:**
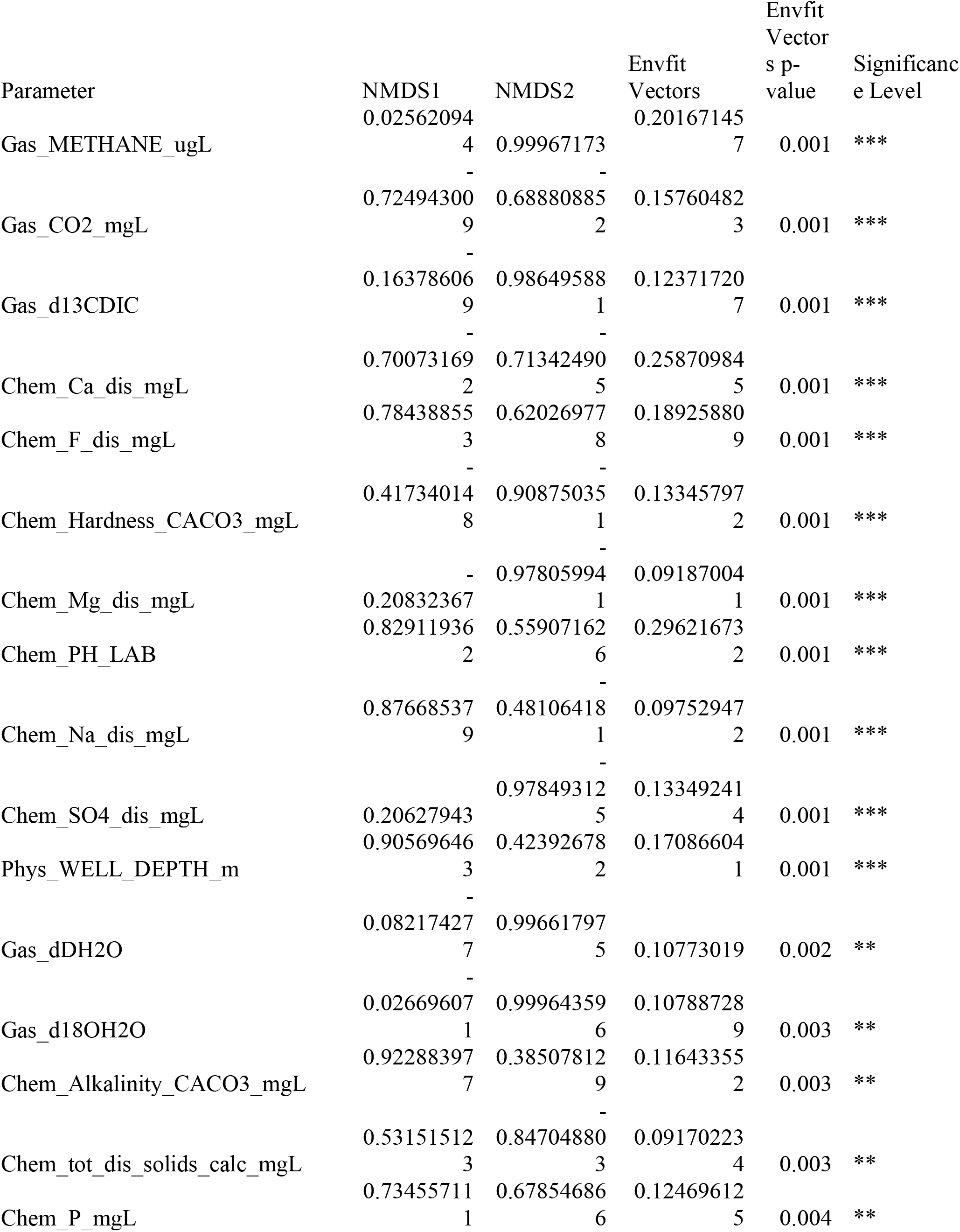
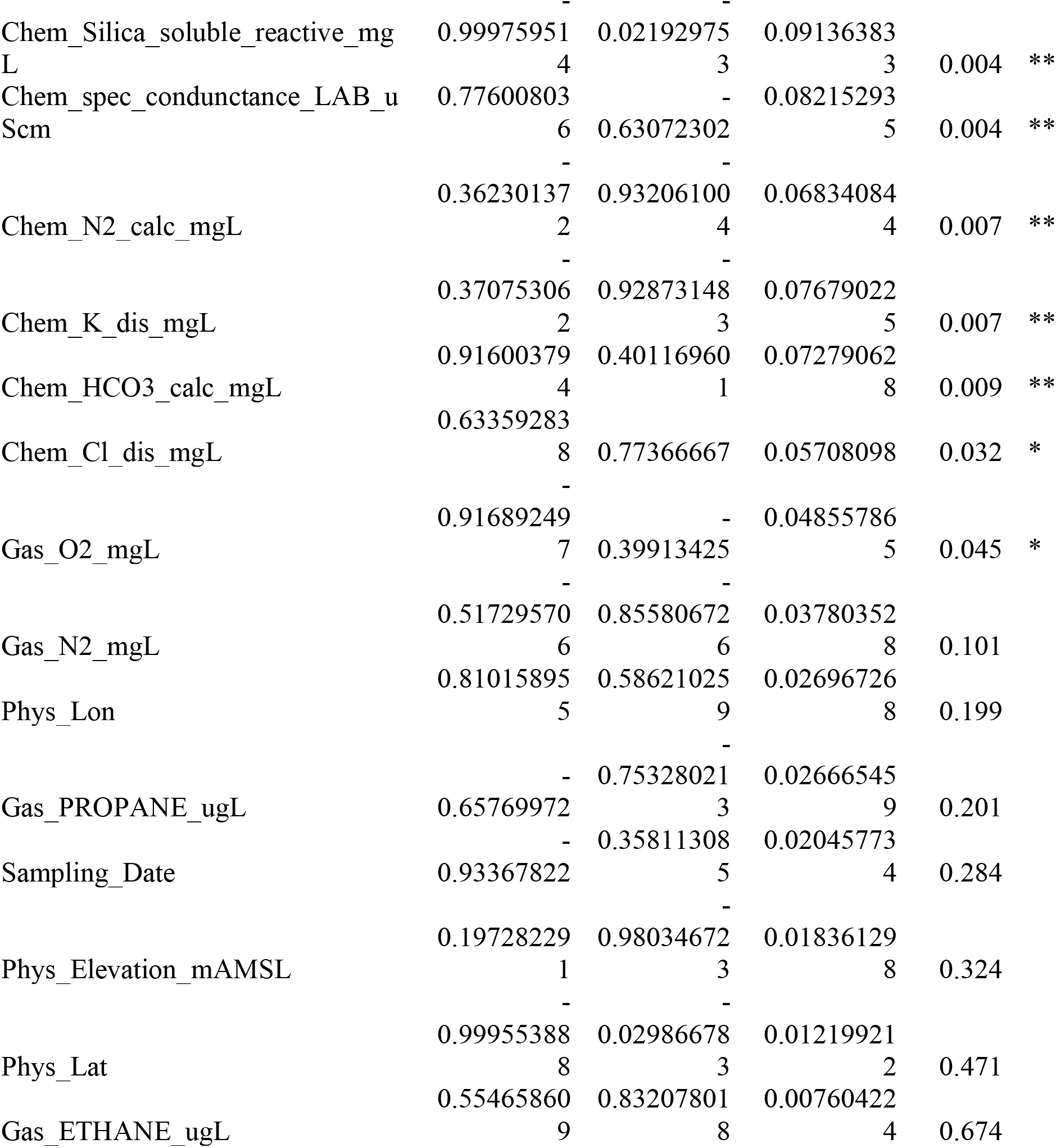
Environmental fitting using bacterial ASV data.

